# Deep learning-enabled design of synthetic orthologs of a signaling protein

**DOI:** 10.1101/2022.12.21.521443

**Authors:** Xinran Lian, Niksa Praljak, Subu K. Subramanian, Sarah Wasinger, Rama Ranganathan, Andrew L. Ferguson

## Abstract

Evolution-based deep generative models represent an exciting direction in understanding and designing proteins. An open question is whether such models can represent the constraints underlying specialized functions that are necessary for organismal fitness in specific biological contexts. Here, we examine the ability of three different models to produce synthetic versions of SH3 domains that can support function in a yeast stress signaling pathway. Using a select-seq assay, we show that one form of a variational autoencoder (VAE) recapitulates the functional characteristics of natural SH3 domains and classifies fungal SH3 homologs hierarchically by function and phylogeny. Locality in the latent space of the model predicts and extends the function of natural orthologs and exposes amino acid constraints distributed near and far from the SH3 ligand-binding site. The ability of deep generative models to specify orthologous function *in vivo* opens new avenues for probing and engineering protein function in specific cellular environments.

## Introduction

An emerging approach for understanding and designing synthetic proteins is learning the design principles of natural proteins evolved through variation and natural selection. These principles are encoded within ensembles of homologous amino acid sequences and define the mapping from primary sequence to multifaceted protein phenotypes, including foldability, biochemical activities, and organismal fitness in a natural biological context [1, 2, 3, 4, 5]. Evolution-based algorithms that learn these rules have the potential to generate new hypotheses for protein mechanism, and to permit the design of diverse synthetic variants with novel functions, with powerful implications for medicine, biotechnology, chemical engineering, and public health [6].

Historically, protein design typically involve physics-based scoring functions that adopt tertiary structure as the central object to bridge sequence to function [7, 8, 9] or involve directed evolution to learn a sequence to function mapping through iterative rounds of mutation and functional selection [10, 11, 12]. In recent years, advances in deep machine learning have driven exciting developments in machine learning-assisted directed evolution (MLDE) [6, 13, 14, 15, 16, 17] that train models to learn the sequence to function map. The central idea of these strategies is to replace a blind mutational search through the vast gulf of protein sequence space with a model-guided search, and to eliminate the need for the direct use of structural information by implicitly representing the underlying physics in the model-learned parameters. The learned models provide a new understanding of the organizing principles of natural proteins at both in terms of general “linguistic rules” underpinning the patterns amino acids in all natural proteins and the local and global epistatic interactions between amino acids in individual proteins that provide for protein phenotypes [18, 19, 5, 20, 21, 22, 23, 24, 25].

Two MLDE approaches that have demonstrated particular promise are direct coupling analysis (DCA) and deep generative modeling (DGM). The essence of DCA is to start with a multiple sequence alignment (MSA) of a protein family and infer a generative model representing the intrinsic constraints on amino acids (the “one-body” terms) and the pairwise interactions between amino acids (the “two-body” terms) [20, 26, 21, 24, 27]. For the cho-rismate mutase enzyme family, recent work showed that the DCA model is sufficient to design of synthetic variants that function in a manner equivalent to natural enzymes both *in vitro* and *in vivo*, in *E. coli* cells [5]. The relative simplicity of the constraints imposed by the DCA model led to considerable sequence divergence in the synthetic proteins, demonstrating access to an enormous space of functional proteins consistent with the evolutionary constraints.

The DCA model is relatively simple because it is inferred only from the first-and second-order statistics of sequence alignments. Given this, it is impressive that it can suffice to capture the design constraints for specifying proteins that can fold and function in their natural cellular context. However, it is also true that the chorismate mutases largely represent a family of orthologs - extant proteins that are descended by speciation events and are expected to share the same function across species. Indeed, a large fraction of homologous chorismate mutases operate in *E. coli* in the specific experimental conditions in which the design was carried out [5]. Such consistency of function in a protein family likely represents a simpler problem for inference of generative models. A deeper and more general test of evolution-based generative models would come from a study of a family of paralogs - proteins that arose through gene duplication events and typically have diverged to carry out distinct and specialized functions. Indeed, paralogs of a protein family are thought to under strong selection to be functionally orthogonal with respect to each other [28], a strategy to ensure specificity in signaling [28, 29] and metabolic [30] pathways. These observations raise the question of whether it is even possible to make generative models for specific orthologs given input data comprising the full spectrum of functional divergences in most protein families.

An ideal model system to investigate this question is the Src homology 3 (SH3) family of protein interaction modules. SH3 domains are small all-beta folds that bind to type II poly-proline containing peptides of the form N-R/KXXPXXP-C or N-XPXXPXR/K-C [31](Fig. 1A) and mediate diverse signaling functions in cells [32]. For example, a C-terminal SH3 domain in the Sho1 transmembrane receptor in fungi (Sho1^SH3^) mediates the response to external osmotic stress through binding to a polyproline ligand in the Pbs2 MAP kinase (Fig. 1B). The Sho1 pathway has been conserved within the fungal kingdom through many speciation events, creating a diverse ensemble of extant Sho1^SH3^ ortholog sequences. In addition, duplication events have occurred during natural evolution, creating many paralogous SH3 domains that have diverged to acquire distinct and non-overlapping ligand specificities. For example, in *S. cerevisiae*, the Sho1^SH3^ is the only SH3 domain amongst 26 other paralogous domains in genome that can support osmosensing in the Sho1 pathway [28]. This exclusivity *in vivo* is recapitulated in direct binding assays with the Pbs2 ligand, demonstrating that the specificity is directly encoded in the Sho1^SH3^ amino acid sequence. For all these reasons, the SH3 domain provides a powerful system to test the generative power of evolution-based models.

**Figure 1:**
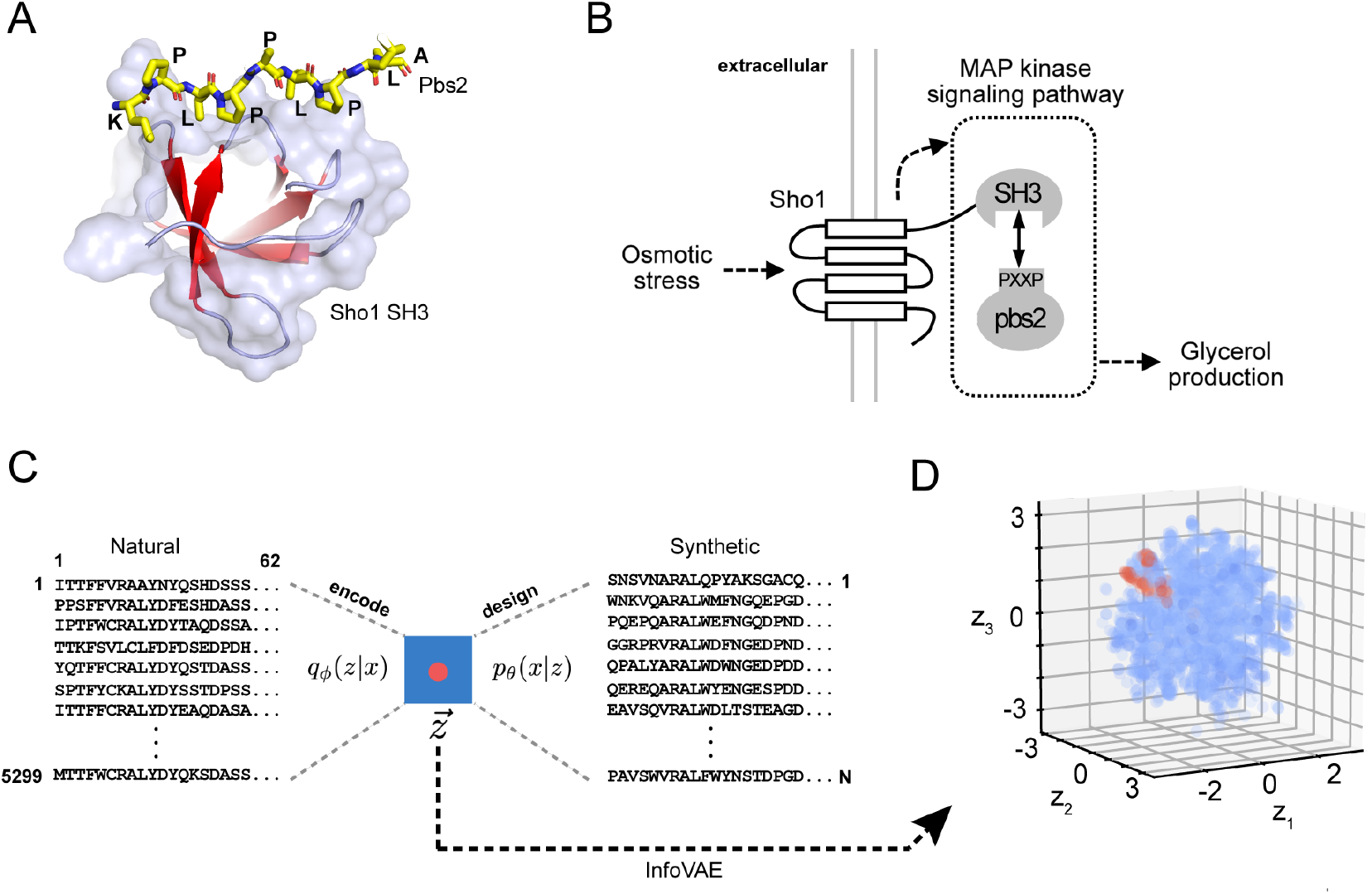
yEvolutionary-based deep generative models of SH3 domains in the context of the yeast high-osmolarity pathway. (A) A structure of the *S. cerevisiae* Sho1^SH3^ domain (PDB 2VKN) in complex with the Pbs2 peptide ligand (yellow stick bonds). SH3 domains are protein interaction modules that bind to polyproline containing target ligands. (B) Binding between the Sho1^SH3^ domain and its target sequence in the Pbs2 MAP kinase kinase mediates responses to fluctuations in external osmotic pressure by controlling the production of internal osmolytes. (C) Schematic of evolutionary-based data-driven generative models, consisting of a compression step (the encoder) that maps a sequence alignment of natural homologs to a low-dimensional Gaussian latent space (blue box), defined by vector 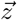 for each sequence, and a decoder which converts latent space coordinates to protein sequences. By definition a VAE is trained to reproduce its inputs; thus decoded sequences represent hypotheses for synthetic members of the protein family. (D) The three-dimensional latent space for the SH3 MSA; the Sho1^SH3^ ortholog group is highlighted in red.

Here, we examine the ability of three modern machine-learning approaches to design of ‘‘synthetic orthologs” of Sho1^SH3^ starting from sequences comprising the full SH3 family. By synthetic orthologs, we mean a designed proteins that span the same diversity as natural Sho1^SH3^ orthologs but that are functionally indistinguishable, both in *vitro* and in vivo. We show that one method (InfoVAE [33]) learns a low-dimensional ‘latent” space that hierarchically organizes SH3 homologs by function and phylogeny. Furthermore, we show that locality in the latent space is both necessary and sufficient to design synthetic Sho1^SH3^ orthologs that bind Pbs2 and support osmosensing in *S. cerevisiae*. Interestingly, constraints on orthology are spread both near and far from the SH3 binding pocket, including many unconserved, solvent-exposed regions that would not be conventionally obvious. The capacity to learn the rules for ortholog function from a functionally diverse protein family provides a platform for a deeper understanding of protein function in a natural biological context.

## Results and Discussion

### Evolution-based deep generative models

We began by constructing a multiple sequence alignment (MSA, see Supplementary Material) of 5299 SH3 homologs, including 3647 fungal domains and 1652 non-fungal domains. The alignment includes all 27 unique paralog groups found in fungal species (from > 150 genomes), representing a deep sampling of the evolutionary record of the fungal kingdom. Sho1^SH3^ orthologs were annotated by fusion to the transmemnbrane portions of the Sho1 receptor rather than by direct alignment scores; thus detection of orthology is independent of sequence similarity within the SH3 domain. This MSA comprises the input data to algorithms that compress the information contained within the natural sequences into a low-dimensional model (Fig. 1C). If the compression captures the essential constraints on folding and binding specificity, it should be possible to design diverse synthetic orthologs of SH3 domains (e.g. Sho1^SH3^) that reproduce the activity and diversity of natural orthologs (Fig. 1C).

The first model we consider is the Boltzmann machine direct-coupling analysis (bmDCA) [26]. The DCA approach assumes that the probability of each natural amino acid sequence *x* = (*x*_1_,…, *x_L_*) to occur is exponentially related to an “energy” function parameterized by the intrinsic constraints on each amino acid *x_i_* at each position *i* (*h_i_*(*x_i_*)) and the pairwise couplings between amino acids (*x_i_,x_j_*) at positions (*i,j*) (*J_ij_*(*x_i_*,*x_j_*)):

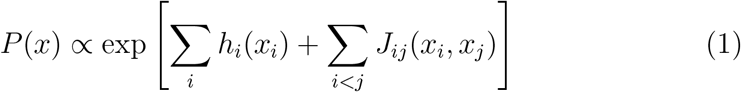

The parameters (*h*, *J*) are trained to reproduce the empirical positional frequencies and pairwise correlations of amino acids (the one- and two-body statistics) in the input MSA. If the model accounts for the information content of natural sequences, synthetic sequences drawn from this probability distribution with low energy (that is, high probability) should be natural-like proteins. Boltzmann machine learning is computationally intensive but provides accurate fitting; for example, the trained bmDCA model for the SH3 family shows excellent reproduction of the input sequence statistics (Fig. S5A). As with any machine learning algorithm, bmDCA involves setting various parameters during model training. Here we follow the approach in previous work [5] to test whether the design of members of the ortholog family studied in that work generalizes to a functionally diverse family of paralogs.

The second class of models we examined are DGMs known as a variational autoencoders (VAEs) [34], consisting of two back-to-back deep neural networks: an encoder *q_ϕ_*(*z*|*x*) that compresses the information content of sequences *x* in the MSA into low-dimensional latent space vectors *z*, and a decoder *p_θ_*(*x*|*z*) that performs the reverse process, transforming latent vectors *z* back into protein sequences *x* (Fig. S1A). If the learning was effective, the latent space should reveal functional and/or evolutionary relationships between sequences, and the decoding process should generate novel sequences from latent space coordinates not occupied by natural sequences. The former operation can be thought of as an interpretive function of the VAE, while the latter represents novel design. In contrast to bmDCA, which learns on the one- and two-body amino acid statistics, the VAE models are trained to reconstruct all features of the input data, and make no assumptions about the form of the sequence-function model. This approach takes advantage of the powerful representational capacity of the deep neural networks [35, 36], and provides a direct solution for designing novel sequences from the latent space without the need for computationally expensive numerical simulations [37, 38, 39, 40].

We implemented two forms of a VAE: (1) a generic, widely-used form that we call the “vanilla-VAE”, and (2) a variant known as an information maximizing VAE (InfoVAE) [33]. While the generic algorithms have proven useful for studying protein properties [41, 42, 43, 44, 45, 25, 37, 39, 38], they can also lead to inaccurate latent inference and non-optimal decoder performance [46, 47]. The InfoVAE addresses these problems, incorporating additional constraints during training models that encourages more accurate decoding from the latent space for design [33]. We present data on both VAE architectures in this work, but for brevity, we illustrate features of the latent space representations in figures below using the infoVAE method.

### The VAE latent space for the SH3 family

Fig. 2 shows the structure of the infoVAE latent space for the SH3 family. A statistical cross-validation approach determines the number of model dimensions; for the SH3 MSA, this indicates a three-dimensional space into which natural sequences are embedded (Fig. 1D). Interestingly, annotation shows that phylogeny is not the primary organizing principle [25]. For example, SH3 sequences from the Saccaromycotina family, the Pezizomycotina class, and the Basidiomycota division are distributed throughout the latent space with no immediately obvious pattern of localization (Fig. 2A). In contrast, sequences are more distinctly organized by paralog group in the fungal genomes. The (Bzz1_1_, Abp1, Rvs167, and Sho1 SH3 domains fall into distinct wedge-like divisions of the latent space (Fig. 2B, S1B, and see Supplementary Information for other paralog groups). However, within each paralog wedge, a sub-organization by phylogeny is evident. For example, for the Sho1^SH3^ group, the Ascomycota and Basidomycota divisions form two branches extending radially from the origin of the latent space, and the non-dikarya SH3 domains are more proximal (Fig. 2B, S2). The precise meaning of the spatial distribution within the patterns is a matter for further study, but we can conclude that the InfoVAE produces a hierarchical organization of SH3 homologs in which functional distinctions are primary, and phylogeny is secondary. In supplementary inforamtion, we show that the vanilla VAE latent space shows a similar hierarchical clustering (Fig. S3).

**Figure 2:**
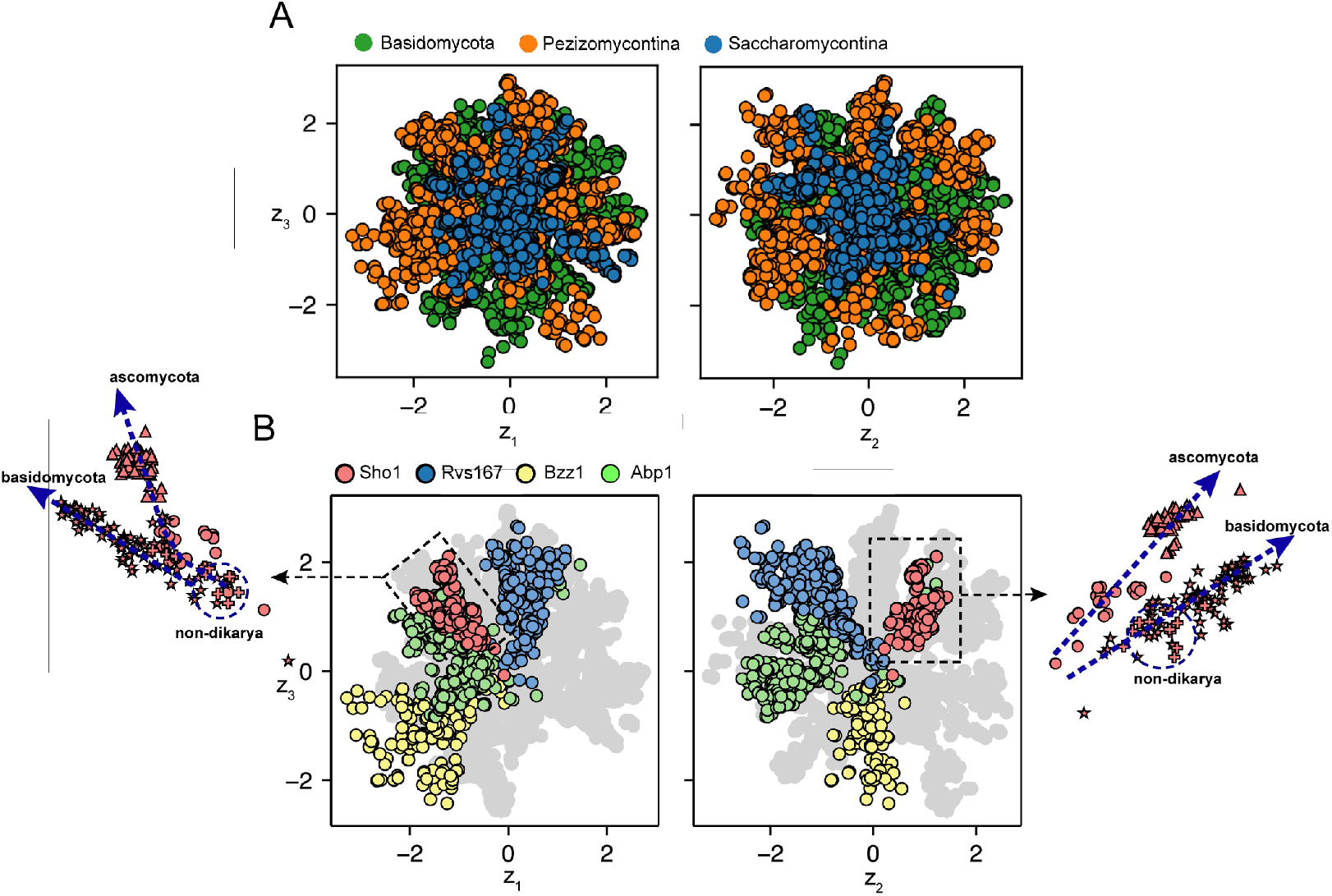
The InfoVAE latents learns a nested hierarchical partitioning of natural fungal SH3 homologs by function and phylogeny. (A) InfoVAE 3D latent space embedding of the 5299 natural SH3 homologs annotated by the three main fungal phylogeny groups. (B) Annotation by paralog group and phylogenetic annotation within the Sho1 paralog cluster (red): Saccaromycotina (circle), Pezizomycotina (triangle), Basidiomycota (star) and non-dikarya (plus). Analogous plots for the remaining paralog groups are presented in Figs. S1 and S2.

To understand how sequences made with just first- and second-order statistics are repersented, we used the trained encoder to embed the bmDCA generated sequences into the latent space (Fig. S5C). The data show that these sequences localize closer to the origin of the VAE latent space, with no observed probability density in the peripheral regions that best distinguish the fungal paralog groups. Note that the VAEs are trained to produce latent space that are multi-dimensional Gaussians; thus, the basic result here is that bmDCA sequences tend towards the average position in latent space. In contrast, VAE sequences extend to more unique positions in the tails of the distribution. These findings suggest that the VAE is learning a different and potentially deeper representation of the information content of SH3 sequences.

### Deep conservation of Sho1 SH3 function in fungal genomes

The localization of fungal ortholog groups in the VAE latent space is consistent with the idea that orthology corresponds to functional similarity [25]. But to what extent do we expect orthologs from diverse species to work in the context of specific model organism under specific experimental conditions? To test this, we developed a high-throughput quantitative select-seq assay for Sho1 pathway function in *S. cerevisiae* (Fig. 3A, and see Methods and Supplementary Material). The assay is based on prior work by Lim and coworkers, who constructed a Sho1 deletion yeast strain in which growth rate can be made to report the binding free energy between the Sho1^SH3^ domain and Pbs2 [28]. Using this strain, we make plasmid libraries in which we replace wild-type Sho1^SH3^ in the Sho1 receptor with natural or synthetic SH3 domains, transform yeast, and grow the entire library in a single flask under selective (1M KCl) conditions for a defined period of time. Deep sequencing of the population before and after selection allows us to compute the enrichment of each allele relative to the wild-type *S. cerevisiae* Sho1^SH3^ (the ‘relative enrichment” or r.e.). Under specific conditions of gene induction, growth time, and temperature, the r.e. quantitatively reports the binding free energy between each SH3 variant and the Pbs2 target ligand (Fig. 3B). The physiological response curve between binding energy and fitness is expectedly sigmoidal, indicating the range of SH3-ligand affinities that can support function *in vivo* under the conditions of these experiments (Fig. 3A). The assay show good reproducibility in independent trials (*ρ*_Pearson_ = 0.87, *n* = 11,442; Fig. S4A) and shows complete dependence on osmosensing (no correlation between selective (1M KCl) and non-selective (0M KCl) conditions (*ρ*_Pearson_ = 0.10, *n* = 10,448; Fig. S4B). Thus, the assay provides a rigorous basis to study large numbers of natural and artificial sequences for *in vivo* functional activity.

**Figure 3:**
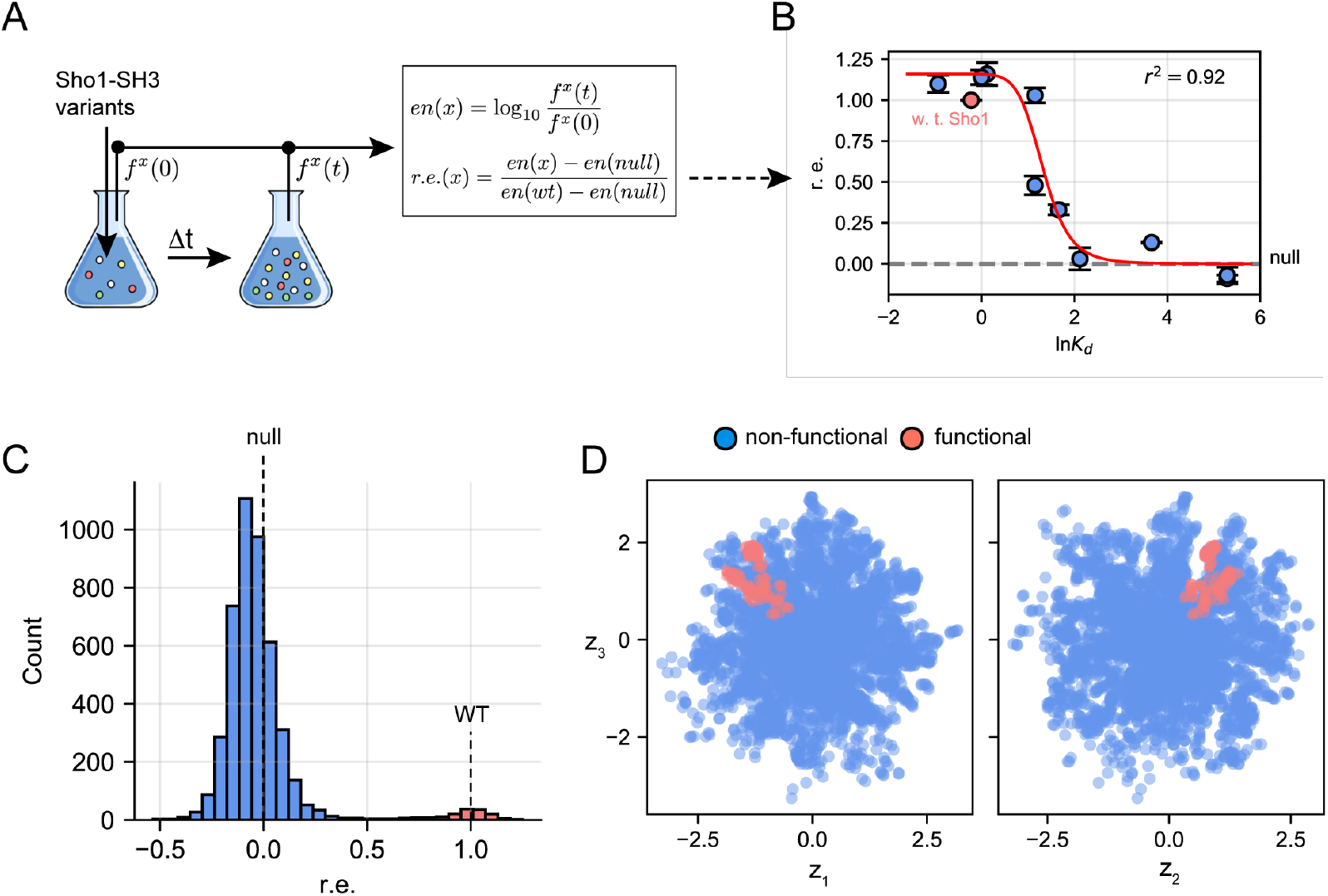
High-throughput select-seq assay for Sho1^SH3^ function in *S. cerevisiae*. (A) Workflow for characterization of yeast high-osmolarity response (i.e., Sho1 functionality). Sho1-deficient *S. cerevisiae* cells (ss101) carrying libraries of variants were grown under selective conditions in 1M KCl media, after which we performed deep sequencing of input and selected population calculation of relative enrichment (r.e.) of each variant. (B) Standard curve linking *in vivo* r.e. with relative binding dissociation constant *K_d_* of pbs2 MAPKK ligand for the Sho1 wild type and a set of 10 synthetic variants with a diversity of *K_d_* values. (C) Observed bimodal distribution of r.e. scores within 1M KCl media of the 5299 natural SH3 homologs. A subset of 132 natural sequences rescue *in vivo* osmosensing function in *S. cerevisiae* (red), which were used for local sampling in VAEs, and the remaining 5167 sequences (blue). (D) Projection of the 5299 natural SH3 sequences into the 3D latent space of the InfoVAE show a crisp clustering between the 132 functional sequences (red) and 5167 sequences that fail to rescue (blue). The rescuing sequences are localized in the vicinity of the Sho1^SH3^ paralog group (c.f. Fig. 2B).

Using the select-seq assay, we examined the ability of all 5299 natural SH3 homologs in the MSA to rescue osmosensing function in *S. cerevisiae*. The result is a bimodal distribution of function, with a small mode (comprising 132 sequences) centered at the level of wild-type Sho1^SH3^ (“functional”) and a large mode centered near to the position of the null allele (“non-functional”). Annotation of the functional sequences shows that they are all orthologs of Sho1^SH3^ throughout the fungal kingdom including Sho1^SH3^ domains from distant Basidomycota and even non-Dikarya species. The ability of these distant Sho1^SH3^ orthologs to work in *S. cerevisiae* to a level indistinguishable from the *S. cerevisiae* ortholog demonstrates deep conservation of Sho1^SH3^ function in the fungal kingdom.

A small subset of natural sequences (331, or 6.2%) fall in an intermediate range between the two modes; these sequences is consistent with prior observations that some fraction of paralogous SH3 domains can partially complement the Sho1 deletion phenotype [28]. A deeper analysis of the “partial-rescue” behavior will be presented elsewhere. For the purposes of this work, this comprehensive study of the function of natural SH3 domains in the *S. cerevisiae* Sho1 pathway provides a reference for assessing the performance of the three evolution-based design algorithms tested here. Given that Sho1^SH3^ orthologs localize to a specific wedge in the InfoVAE latent space (Fig. 2B) and that all the fully functional SH3 domains are Sho1^SH3^ orthologs, it follows that coloring the latent space by the r.e. scores reveals nearly the same organization as coloring by orthology (Fig. 2B, 3D).

### Synthetic orthologs of Sho1^SH3^ from deep generative models

The study of natural SH3 domains frames the problem of learning the design rules for specific orthologs. Only 2.5% of the input MSA displays full rescue of osmosensing, but these sequences represent the deep evolutionary history of the fungal kingdom. Thus, a strong test of the power of models trained on the input MSA is the ability to generate synthetic homologs of Sho1^SH3^ with an efficiency, quality, and diversity that matches the input dataset. To test this, we assayed libraries of synthetic SH3 variants designed from the three models (Fig. 4) and tested them together in a single select-seq experiment.

**Figure 4:**
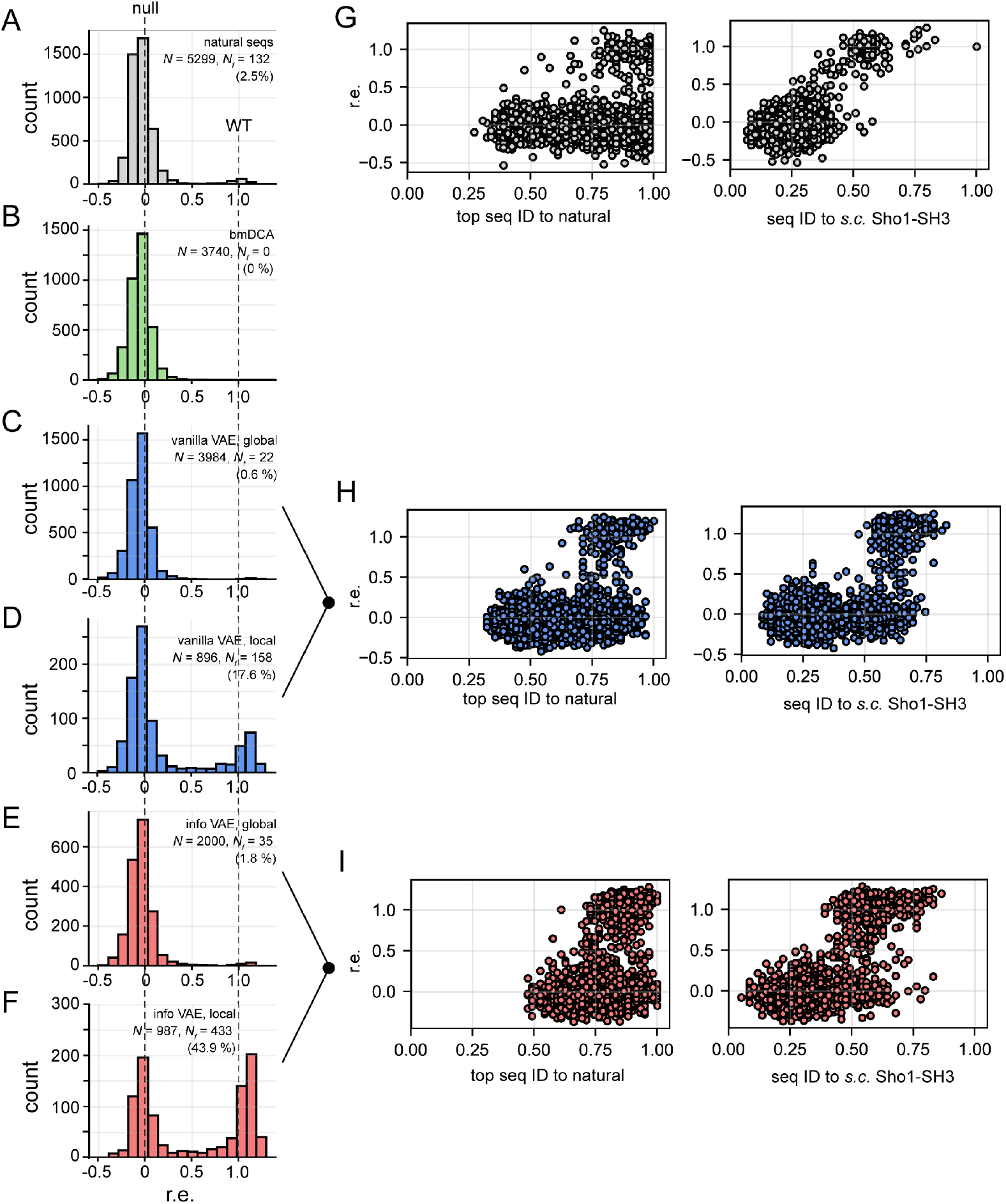
Function and diversity of natural and synthetic SH3 variants. (A-F) Distribution of r.e. scores measured by high-throughput select-seq assay for the 5299 natural SH3 homologs (A), 3740 bmDCA synthetic variants (B), 3984 global (C) and 896 local (D) vanilla VAE synthetic variants, and 2000 global (E) and 987 local (F) InfoVAE synthetic variants. (G-I) Scatterplots of r.e. vs. sequence identity (ID) to the nearest natural homolog or *S. Cerevisiae* Sho1^SH3^ for the 5299 natural sequences (G), 4880 global and local vanilla VAE synthetic sequences (H) and 2987 global and local InfoVAE synthetic sequences (I).

For the bmDCA model, we followed the same protocol in the recent work on the chorismate mutase family [5] to generate synthetic sequences (*N* = 3740) that reproduce the same distribution of statistical energies (e.g. same probability) as the natural homologs (Fig. S5B) [5]. For the SH3 family, the result shows that no bmDCA designed sequences are capable of full complementation of the Sho1 deletion phenotype, though a few sequences fall into a partial rescue range (Fig. 4B). This result is particularly interesting since previous work by Best and colleagues [27] convincingly demonstrates that the bmDCA model is fully capable of producing well-folded and stable SH3 domains. Thus, it appears that bmDCA suffices to make folded SH3 proteins, but at least as tested here, does not capture enough information to specify orthologous function. This outcome could arise either from limitations imposed by using only pairwise statistics in the MSA or from the various approximations and parameter choices used in inferring the model [48]. Regardless, the central conclusion is that at least for Sho1^SH3^, simply reproducing the statistical energies of natural sequences in the bmDCA model is not sufficient to reproduce the distribution of function.

What is the generative capacity of the VAE models? We generated libraries of synthetic sequences from the latent space of both vanilla (N=3984) and infoMAX (N=2000) models by randomly sampling latent space coordinates and passing them through the decoder to convert into protein sequences (Fig. S1A). Re-embedding the designed sequences using the encoder demonstrates that they globally sample the latent space in both models (Fig. S5C). Experimental analysis with the select-seq assay shows that both models are able to produce variants that rescue Sho1 function to the same level as wildtype *S. cerevisiae* Sho1^SH3^ (Fig. 4C, 4E), albeit with different yields. Specifically, 0.6% of vanilla-VAE and 1.75% of infoVAE designed sequences fully function in the Sho1 pathway. A two-sample Kolmogorov-Smirnov test shows that the vanilla-VAE distribution deviates from the natural distribution (*p* = 1 × 10^−4^), but that the InfoVAE distribution is statistically nearly the same (*p* = 0.06). These data show that both VAE models have the capabilities to design functional synthetic orthologs of *S. cerevisiae* Sho1^SH3^ but as expected, the InfoVAE model more accurately represents the design rules embedded in the natural ensemble.

The localization of natural Sho1^SH3^ orthologs in the latent space (Fig. 2B) suggests an additional hypothesis - that sampling in the immediate vicinity of natural orthologs should enrich the yield of synthetic orthologs. To test this, we computed the mean and variance of the functional natural orthologs and designed libraries of sequences from latent space coordinates sampled from the corresponding Gaussian distribution (*N* = 896 and *N* = 987 for vanilla- and info-VAE, respectively). A re-embedding of these sequences shows that they return to the environment from which they were sampled (Figs. 5 and S5C), a quality check on the robustness of the VAE model in these regions. Experimental testing shows that indeed, local sampling produces a much higher density of fully functional synthetic orthologs (Fig. 4D, 4F). An interesting observation is that natural Sho1^SH3^ orthologs fall into phylogenetically defined radially organized sub-regions within an overall space filled out by functional synthetic sequences Fig. 5. Thus, locality in latent space corresponds to locality in function, even for models trained on sequence data alone and no prior knowledge of function.

**Figure 5:**
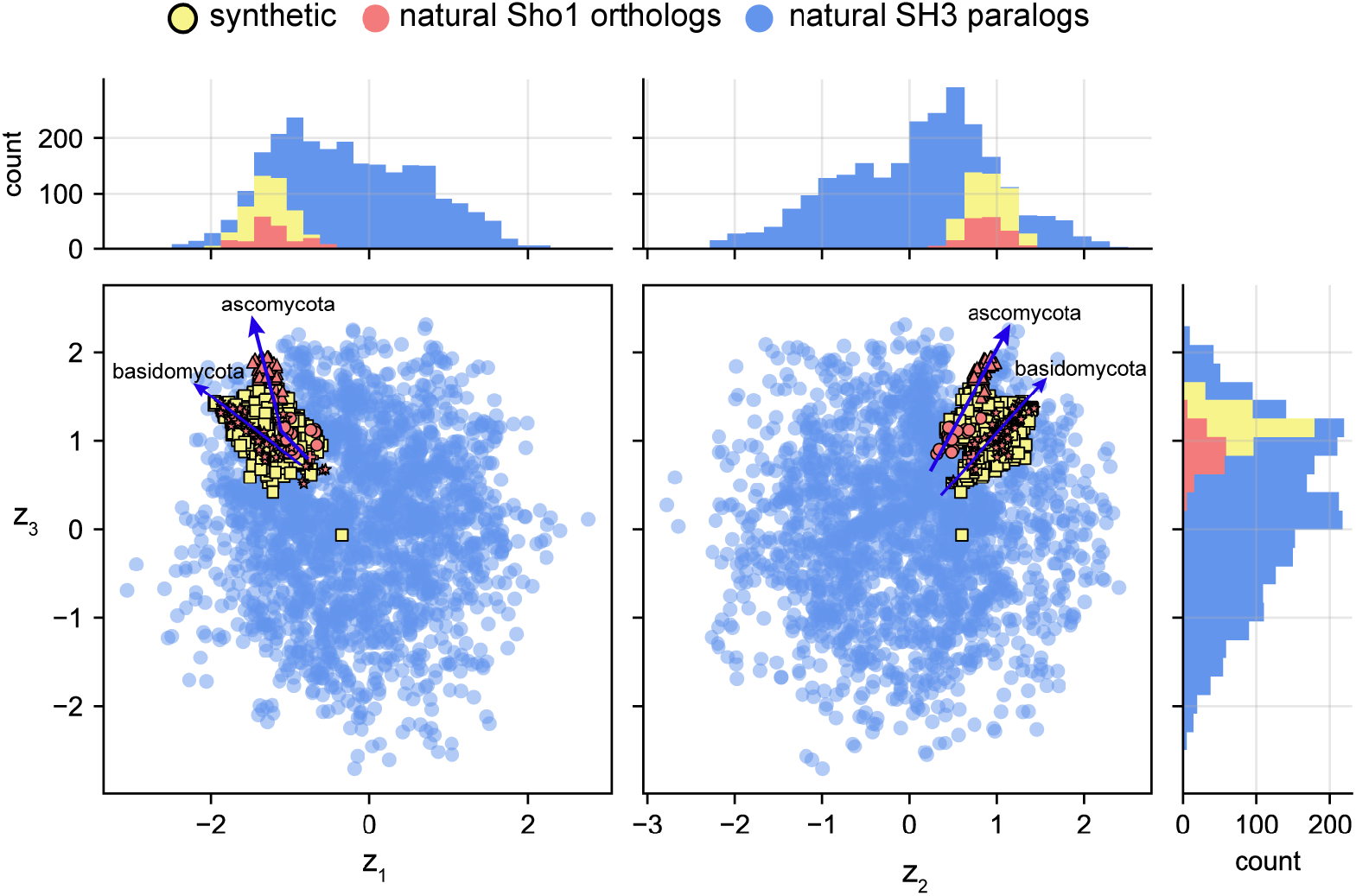
The sequence-function relationship in the infoVAE latent space. Reembedding all synthetic functional sequences in the infoVAE latent space shows that they return to the local environment from which they were sampled, a test of robustness of the model. Natural sequences occupy phylogenetically structures trajectories within an overall wedge-like space that defines Sho1^SH3^-like function.

We selected five synthetic orthologs that show full function *in vivo* for in-depth biochemical characterization. These proteins were expressed in *Escherichia coli* as His6-tagged fusions, purified to homogeneity, and assayed for (1) binding to the *S. cerevisiae* Pbs2 target peptide using a standard tryptophan fluorescence assay [49] and (2) thermal stability by differential scanning calorimetry. The data show that the synthetic proteins are well expressed, soluble, and display a range of binding affinities that are comparable to, or stronger than, the value for wild-type *S.cerevisiae* Sho1^SH3^ (Table 1, Fig. S6). Thermal denaturation experiments show that the synthetic proteins show cooperative unfolding transitions with half-maximal melting temperatures (*T_m_*) and enthalpies of unfolding that span a range around the wild-type protein. Thus, the synthetic variants display biochemical properties similar to natural Sho1^SH3^ domains.

**Table 1:** Biophysial study of five synthetic functional InfoVAE synthetic SH3 variants. ID (WT) = sequence identity to wild-type Shol^SH3^ ([56]), ID (closest) = sequence identity to nearest natural SH3 homolog, *K_d_* = equilibrium dissociation constant for binding the PBS2 target peptide ligand, *T_m_* = half-maximal denaturation temperature (by DCS), Δ*H* = enthalpy of unfolding at the *T_m_*.

| Header | Closest Sho1<sup>SH3</sup> ortholog | ID (WT) | ID (closest) | $K_d$ [μM] | $T_m$ [$^{\circ}$C] | $\Delta H$ [kJ/mol] |
| --- | --- | --- | --- | --- | --- | --- |
| WT | *Saccharomyces cerevisiae* | 1.00 | 1.00 | 3.0±0.1 | 59.1 | 41.2 ± 0.3 |
| InfoVAE_local_1 | *Trichophyton rubrum* | 0.53 | 0.92 | 1.1±0.1 | 44.5 | 41.5 ± 1.9 |
| InfoVAE_local_2 | *Moesziomyces antarcticus* | 0.53 | 0.90 | 0.7±0.1 | 65.0 | 50.9 ± 0.7 |
| InfoVAE_local_6 | *Fistulina hepatica* | 0.54 | 0.83 | 0.3±0.03 | 58.5 | 38.0 ± 0.3 |
| InfoVAE_local_10 | *Trichophyton rubrum* | 0.56 | 0.85 | 2.2±0.4 | 62.5 | 41.6 ± 1.1 |
| InfoVAE_local_11 | *Neurospora crassa* | 0.59 | 0.88 | 0.8±0.04 | 66.5 | 56.3 ± 0.6 |

What is the diversity of the new synthetic variants with respect to natural SH3 domains? For comparison, Fig. 4G shows the distribution of top sequence identities of natural sequences to their nearest natural counterpart or to *S. cerevisiae* Sho1^SH3^. Functional Sho1^SH3^ orthologs are more sequence similar to each other (>60% top-hit identity) than to SH3 paralogs, but can be quite diverged from *S. cerevisiae* Sho1^SH3^ (as low as 40% identity). The vanilla- and info-VAE methods approximate the same diversity, both in terms of distance from all Sho1^SH3^ orthologs and from the *S. cerevisiae* variant (Fig. 4H-I). The ability to reproduce the sequence diversity of natural homologs suggests that the models learn the physical constraints on orthologs without extensive overfitting on irrelevant idiosyncrasies of extant variants.

### Spatial characteristics of Sho1^SH3^ function in the infoVAE latent space

The generative efficiency of the infoVAE latent space inspires a deeper study of how Sho1^SH3^ function maps to latent space position. As noted, the functional natural Sho1^SH3^ and synthetic orthologs are tightly localized to a radially extended wedge-like structure in the VAE latent space (Fig. 5). To make this quantitative, we defined a minimal polygon in the latent space (a so-called “convex hull”) that bounds the natural sequences displaying full function in the *S. cerevisiae* Sho1 pathway (Fig. 6A). The majority of Sho1^SH3^ orthologs in the fungal kingdom (155/172) lie within the hull, and very few sequences within the hull are not functional (Fig. 6B). Also, synthetic orthologs embedding inside the hull show the same distribution of function as their natural counterparts (Fig. 6C-D). Thus, the hull represents a bounding box that defines the space of extant and synthetic functional Sho1^SH3^-like orthologs.

**Figure 6:**
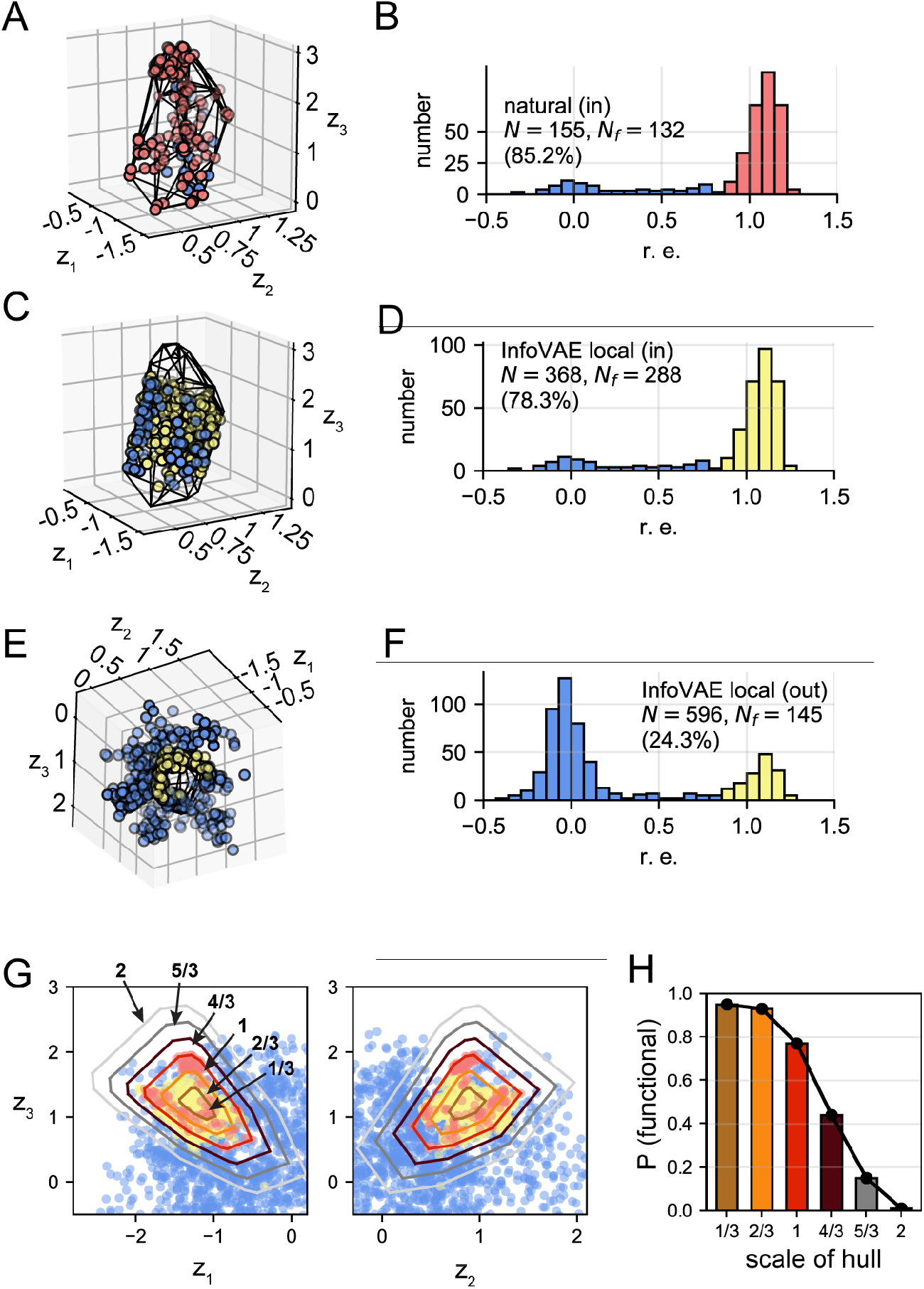
Spatial localization of Sho1^SH3^ function in the VAE latent space. (A-B) Convex hull (black lines) of the natural functional SH3 orthologs (red) defined as the smallest convex polygon that encloses 132 functional SH3 homologs. A small number of 23 non-functional natural sequences (blue) are contained within the convex hull construction. The preponderance 85.2% of sequences contained within the convex hull are functional, indicating that localization within the region of latent space defined by the convex hull is a good proxy for osmosensing function. (C-D) Analysis of the synthetic sequences locally designed by the InfoVAE lying *within* the natural convex hull reveals 288 functional (yellow) and 80 non-functional (blue) synthetic variants, indicating that 78.3% of synthetic InfoVAE variants residing within the convex hull are functional. (E-F) Analysis of locally designed InfoVAE synthetic sequences lying *outside* the natural convex hull reveals 145 functional (yellow) and 451 non-functional (blue) synthetic variants, indicating that 24.3% of local InfoVAE variants residing in the vicinity of the convex hull are functional. (G) Illustration of the hulls scaled by 1/3, 2/3, 1, 4/3, 5/3, and 2 within 2D projections of the InfoVAE latent space and superposed upon the 132 functional natural SH3 orthologs (red), 468 functional synthetic proteins, and the rest of non-functional synthetic proteins (blue) generated by the InfoVAE. (H) Probability (P) of functional natural and InfoVAE designed sequences contained within each hull as a function of scaling factor.

How does Sho1^SH3^-like function change as one exits the convex hull? Consistent with the idea that the hull defines Sho1^SH3^ function, synthetic orthologs re-embedding outside the convex hull are largely non-functional, with the few that do show Sho1^SH3^-like function occurring in the immediate shell outside the hull (Fig. 6E-F). To quantitatively examine how Sho1^SH3^ function varies across the boundary of the hull, we computed the probability of functional sequences in the *S. cerevisiae* Sho1 pathway as a function of scaled volume shells of the convex hull moving from within the hull to outside (Fig. 6G-H). The data show that Sho1^SH3^-like function drops sharply across the boundary, supporting the idea that the hull largely encloses the sequence rules for Sho1^SH3^ function.

An interesting feature is that the immediate environment outside the convex hull includes some bonafide Sho1^SH3^ synthetic orthologs (Fig. 6E, yellow symbols). This demonstrates a principle of extrapolation in the VAE model in which the space of designable functional sequences extends beyond the limits defined by natural orthologs alone.

### Locality in the latent space exposes global amino acid constraints

The finding that locality within the convex hull of the InfoVAE latent space defines Sho1^SH3^ function provides an opportunity to examine the pattern of amino acid constraints that specifically underlie orthologous function. A simple approach is to compare the conservation of sequence positions in sequences sampled globally from the VAE latent space with that from sequences embedded within the convex hull (Fig. 7). In essence, this analysis provides as first-order view of where the “extra” constraints to be a Sho1^SH3^ ortholog occur in the amino acid sequence. The conservation pattern for globally sampled sequences is nearly the same as for the natural MSA (Fig. S7), a result consistent with the finding that global design reproduces the distribution of function in the natural MSA. However, it is quite different for sequences sampled within the convex hull bounding Sho1^SH3^-like function (Fig. 7A). The differences in conservation can be modeled by a double Gaussian mixture model, providing a statistical basis to identify positions that contribute the most to Sho1 function (Fig. 7B). The extra constraints for Sho1^SH3^ function arise both at known specificity determining sites in the ligand binding pocket [50, 51] and at a set of weakly-conserved and solvent-exposed positions distributed throughout the protein structure (Fig. 7C). These findings illustrate the use of VAE models to provide new hypotheses for mechanisms of protein function in specific cellular contexts *in vivo*.

**Figure 7:**
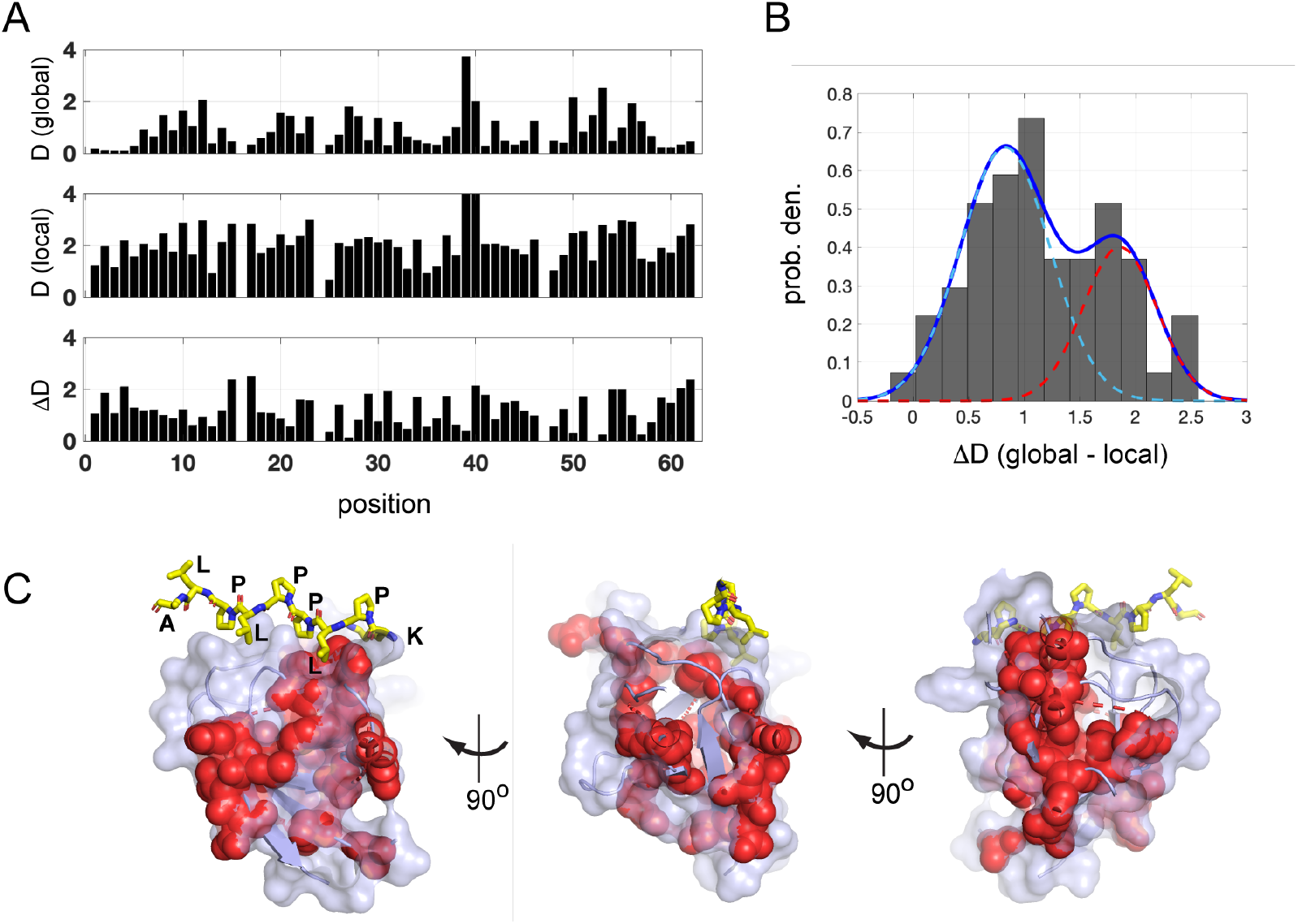
The structural basis for Sho1^SH3^ function. (A) Positional conservation (measured by Kullback-Leibler relative entropy D) in sequences sampled globally from the InfoVAE latent space (top panel),locally from the convex hull bounding functional natural sequences (middle panel), and the difference of the two (bottom panel). This analysis exposes the extra constraints in SH3 domains to be specifically functional in the Sho1 osmosensing pathway. (B) The distribution of differences in conservation, with a fit to a double Gaussian mixture model (blue). For illustrative puproses, the mixture model helps to identify a population of 21 positions showing the largest change in conservation (red curve). (C) The positions showing the largest change in conservation (red speheres) are located at specificity determining regions of the ligand binding pocket and extending throughout the tertiary structure. The imgages show three rotations of the Sho1^SH3^ structure, with the co-crystallized Pbs2 peptide ligand in yellow stick bonds.

## Conclusion

In this work, we show that the latent space of variational encoder models trained on homologs of the SH3 protein family capture the rules for specifying folding and function of specific orthologs of the family. Using this approach, we generated hundreds of sequence-diverse synthetic orthologs of the Sho1^SH3^ domain that support osmosensing in *S. cerevisiae* to an extent comparable to the wild-type domain. This result expands the use of generative models to protein families in which functional diversification leaves only a small fraction of sequences in the input data (< 3%) that can operate in a specific cellular and genome context. In addition, the data show that Sho1^SH3^ function is localized to a small volume of the VAE latent space, and that localization to that volume is nearly necessary and sufficient to specify synthetic orthology. It is interesting that extant natural orthologs occupy only sparse, phylogenetically-structured trajectories within the volume (red symbols and blue arrows, Fig. 2B and Fig. 5). A logical interpretation is that natural sequences are constrained not only by the need to fold and to function, but also by the stochasticity and historical contingencies of natural evolution. Thus, natural sequences are forced to organize into specific sub-regions within a large design space controlled by the underlying selection pressures. In this sense, functional synthetic sequences arising from non-natural regions of latent space may be thought of as alternative histories that could have occurred (but did not) in the history of evolution.

From a practical perspective, these findings suggest that even with no supervision from experimental data, the VAE is distilling the essential physical constraints on folding and function and, at least to some extent, removing pure historical constraints. Thus, the model opens up a vast space of synthetic solutions that span a range of biochemical phenotypes with regard to binding affinity and stability. It may be possible to use the initial round of synthetic design to iteratively train the models to recognize directions in multi-dimensional phenotypic space that deviate from the history of natural selection, but that may be of practical value. Such a semi-supervised design process might represent a practical approach to the design of optimized or even novel phenotypes [52, 37]. From a fundamental point-of-view, the study of iteratively trained models may provide insight about the capacity of natural proteins for phenotypic innovation, a central property of systems evolving under fluctuating conditions of selection [53].

Due to extensive past work documenting tight functional specificity *in vivo* and great functional diversity [28, 51], the SH3 domain family serves as a productive model system for studying the generative potential of data-driven models. However, the choice of the experimental system, algorithms for model construction, and assay technologies are otherwise unremarkable. Thus, we expect the findings here to be of general impact for understanding and engineering diverse protein functions in specific environments. both *in vitro* and *in vivo*.

It is worth noting the conceptual distinction of evolution-based models from the extensive previous work in making models for proteins. All models for function and design represent a attempt to define rules of phenotypic variation by locality in some space of representation. For example, inspired by the steep distance- and geometry-dependence of the fundamental forces between atoms, physics-based design often focuses on local environments of tertiary structure to vary biochemical activities. For example, computational redesign of enzyme function typically involves variation of residues in the immediate contact environment of target ligands [8], a strategy to contain the complexity of the search process. An alternative method - directed evolution - uses rounds of mutagenesis to search locally in the sequence space surrounding a natural protein to design new activities. The logic that evolutionary constraints force the local sequence environment of natural proteins to be densely populated and functionally connected such that it is possible to transit to new phenotypes through paths of single-step variations [54]. Thus, an iterative search of the local environment is a productive approach for discovery of novel functions [55]. The data presented here suggests an alternative principle of design - locality in the latent space of the evolution-based models. This principle does not limit variation to local primary or tertiary structure environments; instead, it is organized by the patterns of epistatic interactions that underlie protein folding and function. Non-linear learning tools such as the VAE are specifically capable of abstracting these complex features of proteins from extant sequence data, and thus open up an enormous new space for protein design. What is perhaps most surprising is the ability of these models to learn generative rules for protein phenotypes from the limited and biased sampling of available sequuences comprising a protein family [48]. The results speak to the relative simplicity of the information stored in natural protein sequences and provide a starting point to understand how basic physical and evolutionary constraints acting on natural proteins.

## Acknowledgements

We gratefully acknowledge support from the Machine Learning in the Chemical Sciences and Engineering program of The Camille and Henry Dreyfus Foundation. This work was supported with funding by the University of Chicago Data Science Institute (DSI). This work was completed in part with resources provided by the University of Chicago Research Computing Center. We gratefully acknowledge computing time on the University of Chicago high-performance GPU-based cyberinfrastructure supported by the National Science Foundation under Grant No. DMR-1828629 and grant NIH RO1GM141697 from the National Institutes of General Medical Sciences (R.R.). We thank members of the Ranganathan and Ferguson groups for helpful comments on manuscript, K. Husain for guidance on highly-efficient yeast transformation, E. Hinds for guidance on growth assays, and B. Andrews for guidance on Illumina MiSeq sequencing.

## Conflict of Interest Disclosure

R.R. and A.L.F. are co-founders and consultants of Evozyne, Inc. and co-authors of US Provisional Patent Application 62/900,420 and International Patent Application PCT/US2020/050466. A.L.F. is also co-author of US Patent Application 16/887,710, US Provisional Patent Applications 62/853,919 and 63/314,898, and International Patent Application PCT/US2020/035206.

