## Supplementary Information for "Deep learning-enabled design of synthetic orthologs of a signaling protein"

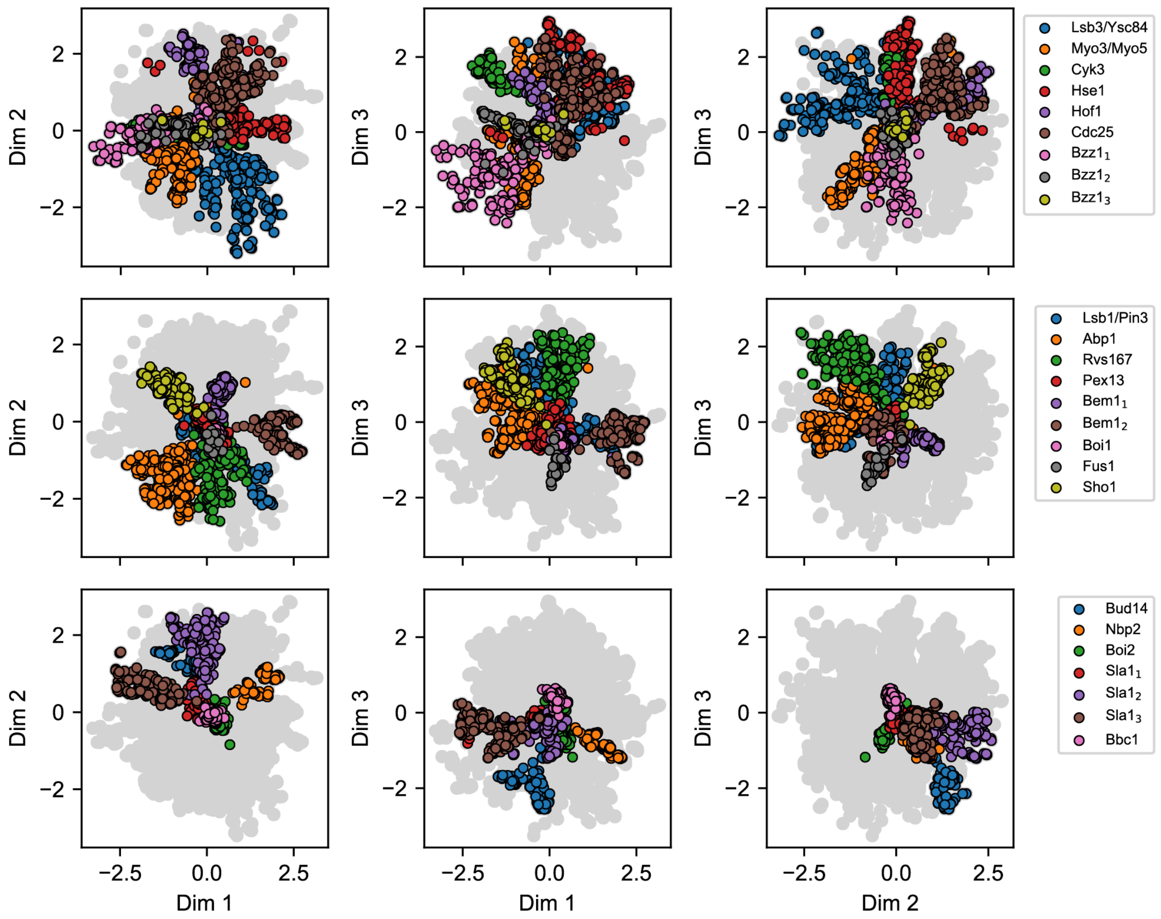


Figure S1: Organization of the 25 fungal SH3 ortholog groups in the 3D InfoVAE latent space. Ortholog groups comprise approximately radially extended wedge-like regions.

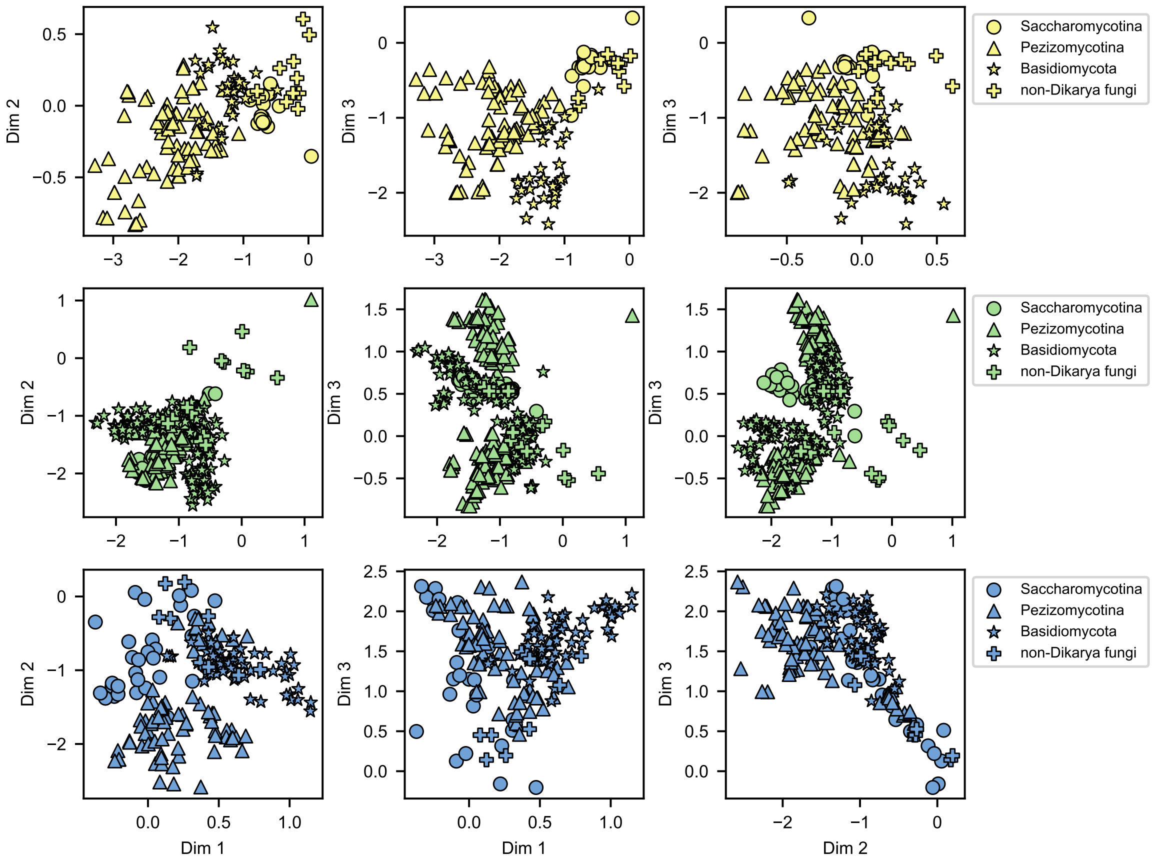


Figure S2: InfoVAE latent space embeddings and phylogenetic annotation of the three additional SH3 paralog groups Bzz11 (yellow), Abp1 (green), and Rvs167 (blue), showing the nested hierarchical organization by function (color) and phylogeny (symbol).

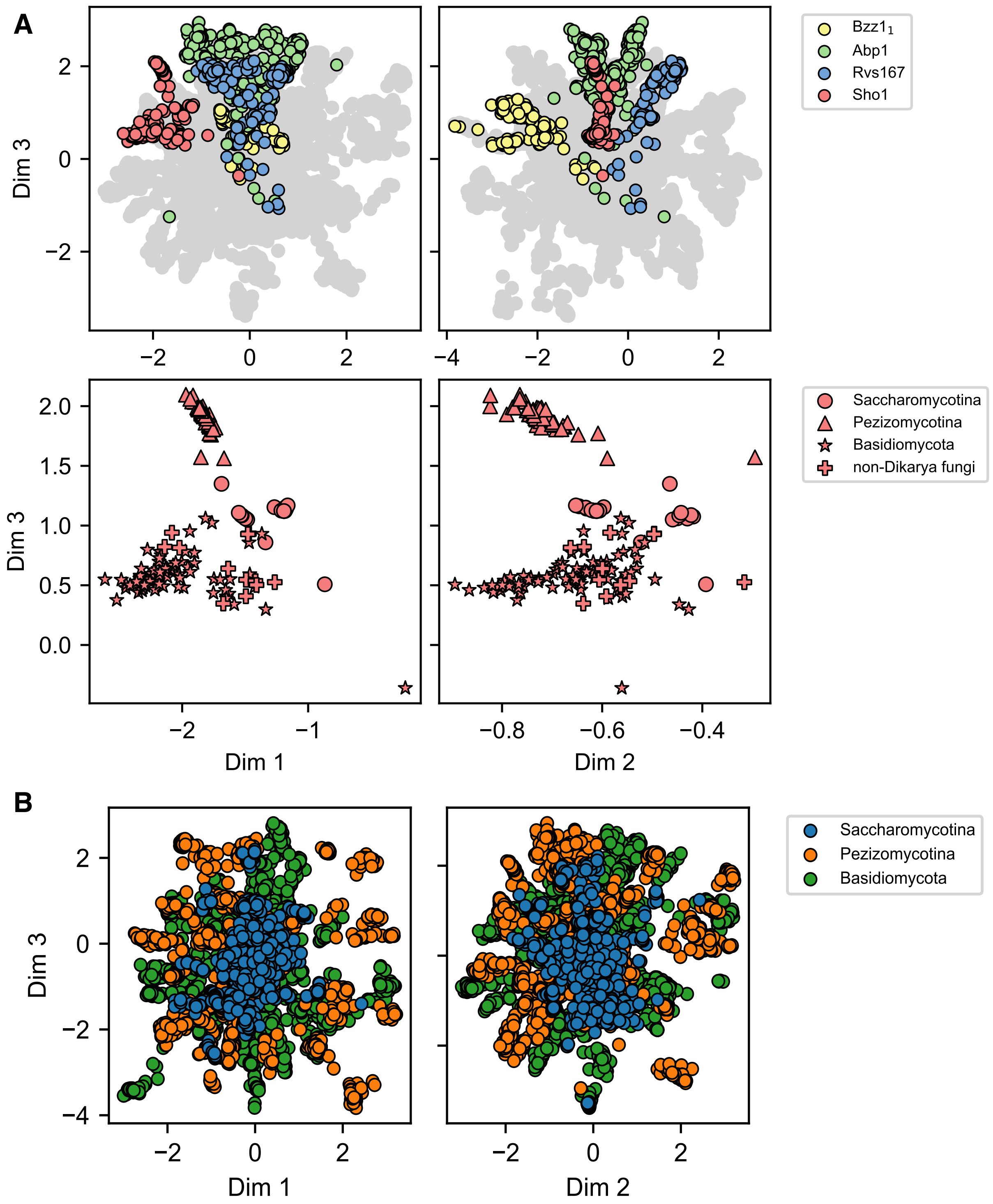


Figure S3: The latent space of the vanilla VAE. (A) Annotation by paralog group and phylogenetic annotation within the Sho1 paralog cluster (red). Similar to the infoVAE (Fig. 2), the vanilla VAE hierarchically organizes by function and phylogeny. (B) The vanilla 3D latent space embedding of the 5299 natural SH3 homologs annotated by the fungal phylogeny groups shown in Fig 2A.

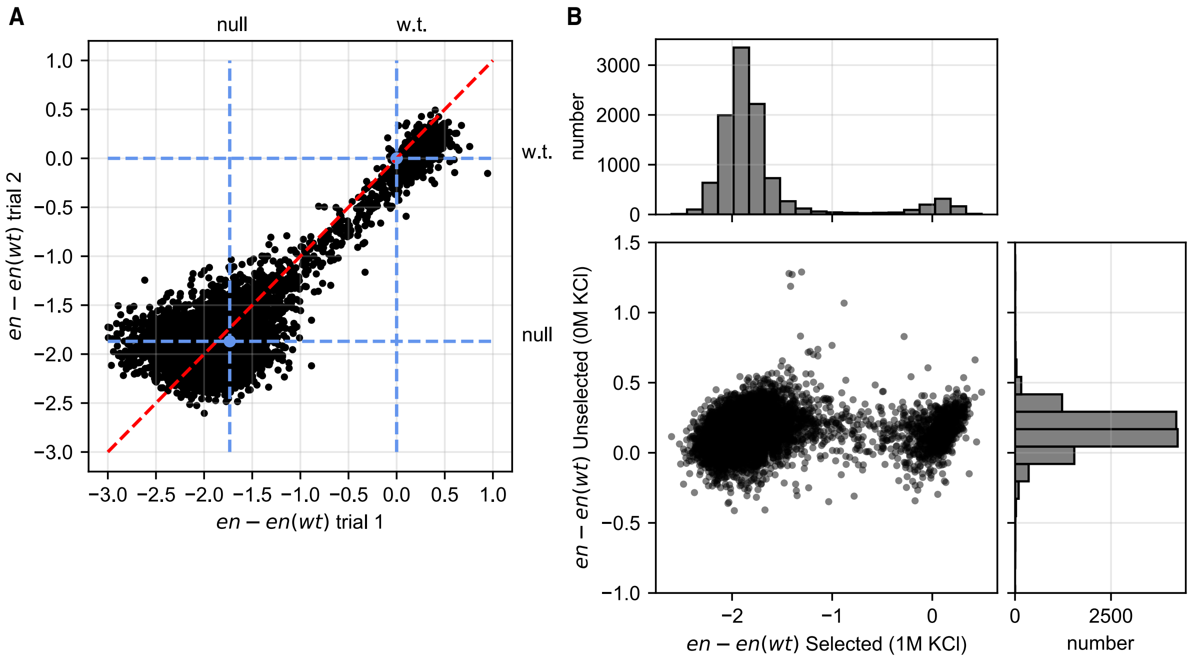


Figure S4: Validation of the high-throughput select-seq assay. (A) A scatterplot of the enrichment score relative to wild-type $[en - en(wt)]$ for two independent ($n$ = 11,442) trials of the select-seq assay under the same experimental conditions. The position of the wild-type Sho1 sequence and the null allele (no Sho1 activity) are indicated by the blue circle and blue dashed lines. The red dashed line is the identity trace. Values at low values of $[en - en(wt)]$ are subject to more variability as expected from poorer counting statistics. The data show that the select-seq assay shows good reproducibility between independent runs ($\rho_{\text{Pearson}} = 0.87$, $n$ = 11,442, $p < 1 \times 10^{-307}$ ). (B) The relationship between $en - en(wt)$ of the SH3 genes in *S. Cerevisiae* grown in selective (1M KCl) and non-selective (0M KCl) media. No statistically significant correlation is observed ($\rho_{\text{Pearson}} = 0.10$, $n$ = 10,448, $p = 6 \times 10^{-23}$). This control experiment shows that bimodal distribution of enrichment in the selected population resulted from differential adaptability under high osmotic pressure conditions.

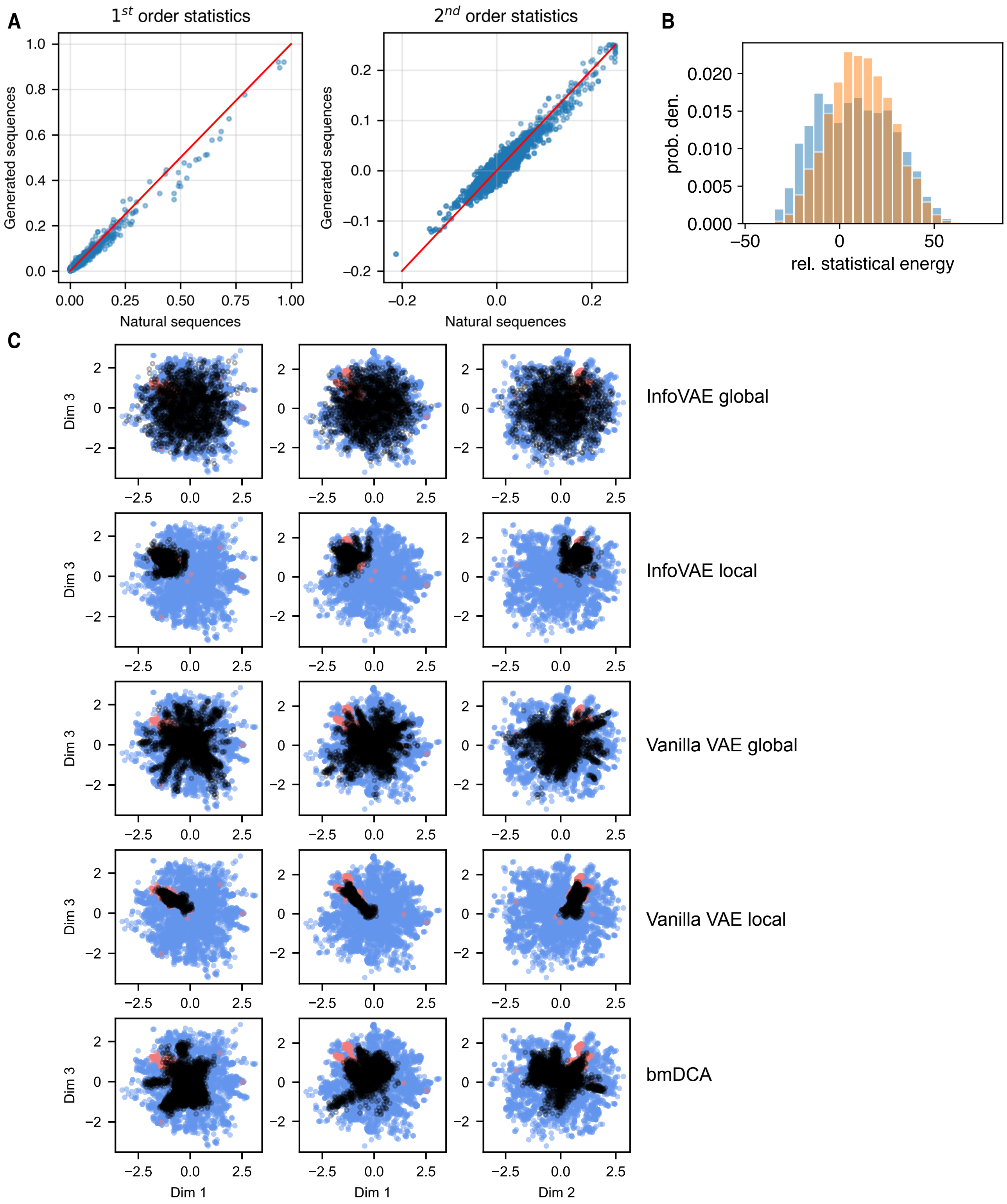

A
$1^{st}$ order statistics
$2^{nd}$ order statistics
Generated sequences
Natural sequences
B
prob. den.
rel. statistical energy
C
Dim 3
Dim 1
Dim 2
InfoVAE global
InfoVAE local
Vanilla VAE global
Vanilla VAE local
bmDCA

Figure S5: Analysis of and sampling from trained generative models. (A) Parity plot of the one- and two-body amino acid frequencies computed over the 5299 natural sequences and an ensemble of 3740 bmDCA designed synthetic variants sampled at $T = 1$. The red lines indicate the identity relationship. The excellent agreement (one-body: $\rho_{\text{Pearson}}$=0.99, $n$= 5000, $p < 1 \times 10^{-307}$; two-body: $\rho_{\text{Pearson}}$=0.82, $n$= 5000, $p < 1 \times 10^{-307}$) demonstrates the validity of the bmDCA model in accurately learning and reproducing the one- and two-body amino acid frequencies. (B) Distribution of statistical energies for natural (blue) and bmDCA designed sequences sampled at $T = 0.9$ (orange). The two distributions lie on the same range and with a similar distribution. (C) Projection into the 3D InfoVAE latent space of the synthetic sequences designed by global ($n$=2000) and local ($n$=987) sampling over the InfoVAE latent space, global ($n$=3984) and local ($n$=896) sampling over the vanilla VAE latent space, and MCMC sampling from the bmDCA generative model ($n$=3740). Designed sequences are shown in black and superposed on the $n$=170 natural SH3 homologs that possess high-r.e. scores and rescue osmosensing function (red) and the remaining $n$=5129 that fail to rescue (blue).

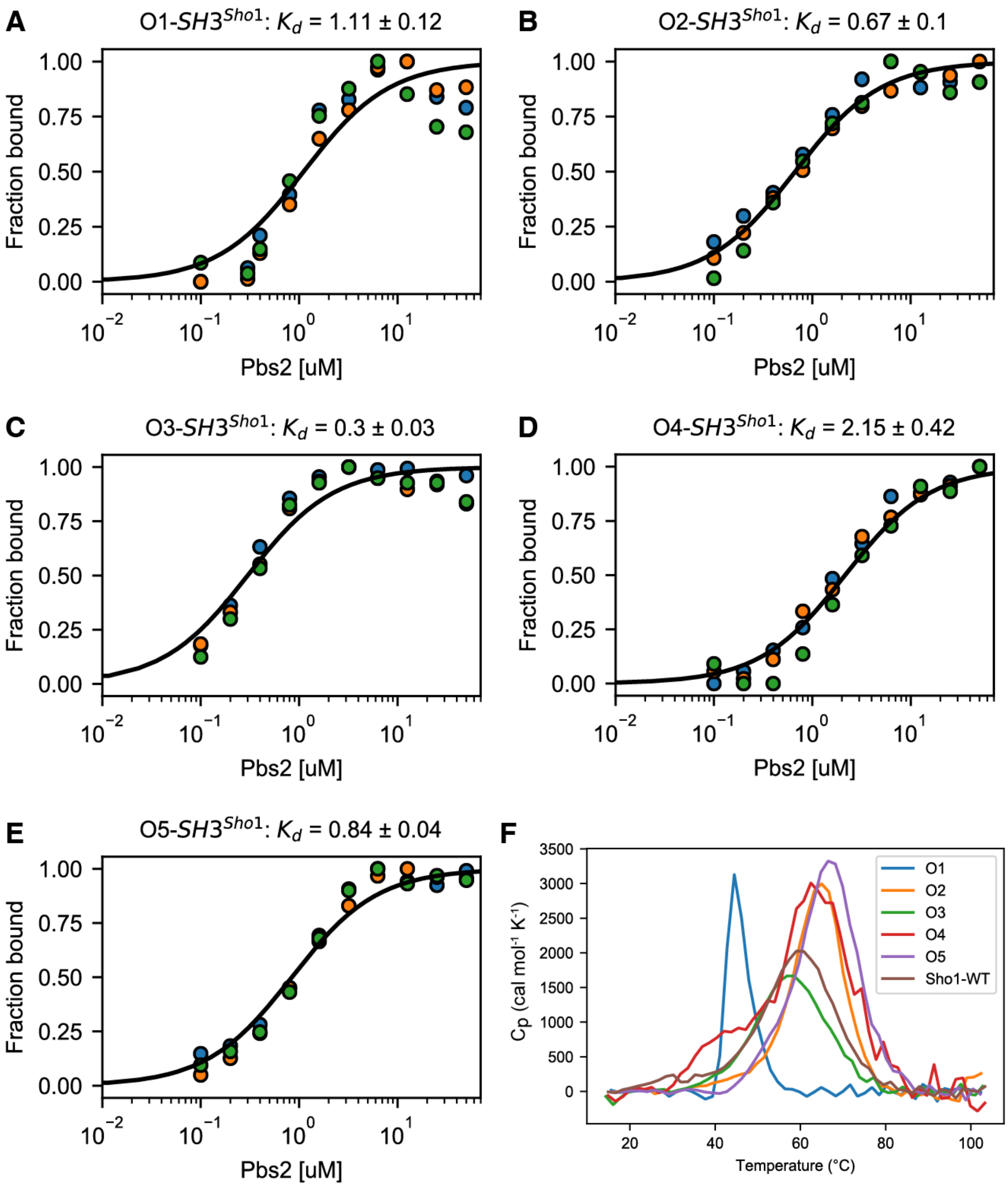


Figure S6: Biophysical analysis of the five designed functional Sho1-SH3 orthologs listed in Table 1. (A-E) Binding isotherms of designed sequences InfoVAE_1 (O1), InfoVAE_2 (O2), InfoVAE_6 (O3), InfoVAE_10 (O4) and InfoVAE_11 (O5) against a PBS2 peptide ligand (see Methods). Colors represent independent titrations. (F) Denaturation curves for the five designed sequences by differential scanning calorimetry (DSC).

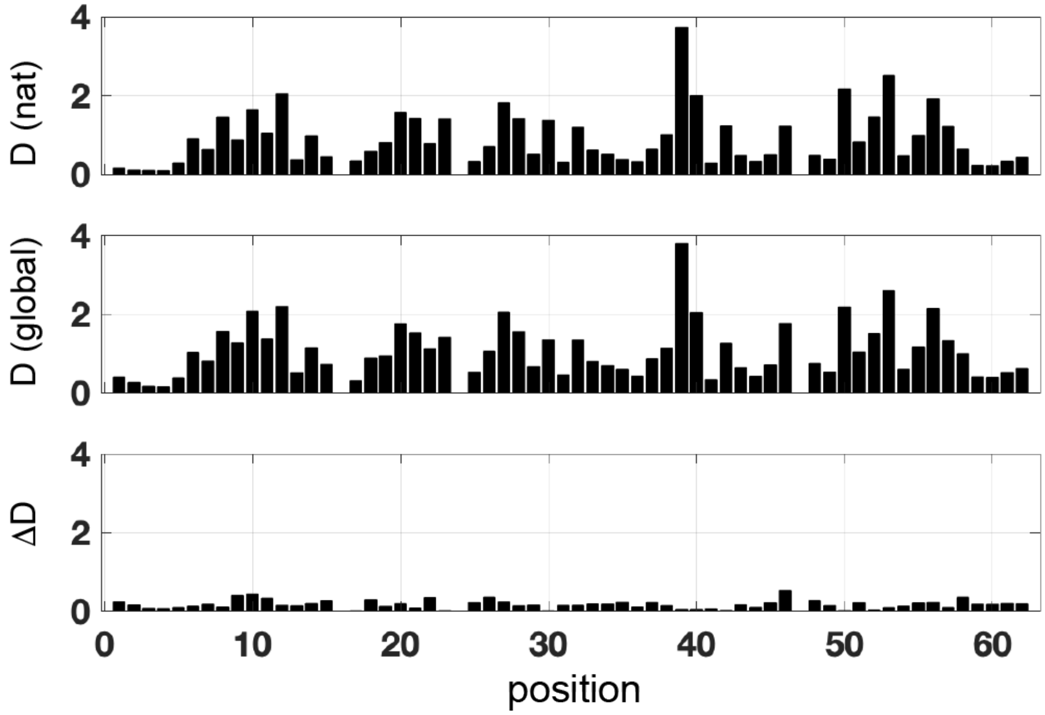


Figure S7: Positional conservation (measured by Kullback-Leibler relative entropy D) in natural SH3 sequences (ope panel), sequences sampled globally from the InfoVAE latent space (middle panel), and the difference of the two (bottom panel). The data show that natural and globally designed sequences display essentially the same pattern of positions conservation.

### Materials and Methods

SH3 protein data collection, preprocessing, and multiple sequence alignment

A total of 7865 sequences of SH3 domains, including 5611 fungal and 2254 non-fungal, were acquired from the JGI (https://jgi.doe.gov), PFAM (https://pfam.xfam.org/) and NCBI (https://www.ncbi.nlm.nih.gov) databases. The sequences were aligned into a multiple sequence alignment (MSA) using PROMALS 3D [1]. The resulting MSA was subject to a final round of manual adjustment using standard rules and trimmed sequentially to retain positions corresponding to residues 2-15, 18-45, and 47-63 of *S. cerevisiae* Sho1$^{SH3}$ (PDB ID 2VKN), to remove sequences with unknown residues, fewer than 56 residues, more than 5% positions trimmed, and duplicates. The final MSA contained 5,299 sequences and 59 positions. For sequences designed from this dataset, the three removed positions in 2VKN were back inserted to make the final length 62 for the purposes of synthetic gene synthesis.

bmDCA inference and sequence generation

The 5299 natural sequences in MSA were used to infer the bmDCA model [2], assigning a probability $P(a_1, ..., a_L) = \frac{1}{Z}\exp\{-H(a_1, ..., a_L)/T\}$ to each aligned sequence $(a_1, ..., a_L)$ with $L = 59$. The statistical energy $H(a_1, ..., a_L) = -\sum_{1\leq i<j\leq L} J_{ij}(a_i, a_j) - \sum_{1\leq i\leq L} h_i(a_i)$ of the Potts model is given in terms of the direct coevolutionary coupling $J_{ij}(a, b)$ between amino acids $a$ and $b$ at positions $i$ and $j$, and propensities $h_i(a)$ for the usage of amino acid $a$ at position $i$. The bmDCA model was inferred at $\lambda = 0.01$ and M=500 using 1600 thermalization steps, and the temperature $T$ is set to unity during inference. The accuracy of the inferred model was checked by comparing first order empirical frequencies $f_i^a$ for each amino acid $a$ at position $i$, and the joint frequencies $f_{i,j}^{ab}$ of amino acids $(a, b)$ at positions $(i, j)$, between the MSA and sequences generated by MCMC at $T = 1$. After the correspondence of the natural and predicted one- and two-body amino acid statistics were validated (Fig. S5A), the final bmDCA designed sequences for experiments were sampled under a lower temperature $T = 0.9$ to produce sequences that have compatible statistical energies with natural sequences (Fig. S5B) [2]. Codes for bmDCA are available at https://github.com/ranganathanlab/bmDCA [3].

Vanilla VAE inference and sequence generation

Each natural homolog within the MSA was converted into a one-hot encoded tensor [4], which maps each individual amino acid label found along a specific sequence into a vector consisting of zeros and ones, where the value 1 indicates the amino acid label. The unique labels consist of at maximum 20 amino acids and deletion gap from the multiple sequence alignment algorithm, and each amino acid position was indexed individually to avoid all-zero features. Thus, the MSA with size $5299 \times 59$ is converted into size $5299 \times 1178$, where the values 5299 and 1178 corresponds to the number of natural homologs used for training and length of the one hot encoded vectors. By using a training dataset that consists of homologs, a variational autoencoder (VAE) was employed as a generative model admitting a low-dimensional embedding of sequence space [5, 6]. With the ability to capture meaningful SH3 evolutionary information through the latent space, this modeling approach has been attractive for protein design, while also introducing the opportunity to capture the distribution of this large homology family which can lead to better understanding of the evolutionary constraints for orthology and paralogy. For example, the decoder can sample from the latent space embedding and generate new artificial and functional protein sequences, while also localizing function within the latent space.

A standard "vanilla" VAE [6–14] was trained to learn the joint probability from Bayes inference: $p_\theta(x, z) = p(z)p_\theta(x|z) = p_\theta(x)p_\theta(z|x)$, where $\theta$ represents learned parameters of the joint distribution, $z \in Z$ represents latent variables, and $x \in X$ represents each sequence $x$ in the training set $X$ of MSA. $p_\theta(x, z)$ denotes the probability of correctly constructing a sequence like those in $X$ given a $z$ from the latent distribution $p(z)$. Each designed sequence $\hat{x}$ was generated by the decoder $p_\theta(\hat{x}|z)$ and $z$ was sampled from $p(z)$. To learn $p_\theta(\hat{x}|z)$, we need to approximate $p_\theta(x)$:

$$p_\theta(x) = \int p_\theta(x|z)p(z)dz \tag{S1}$$

Because it is intractable to directly compute parameters $\theta$ for the probability $p_\theta(x)$, we applied an approximation method called variational inference. Namely, we trained the encoder $q_\phi(z|x) \sim \mathcal{N}(z|\mu_\phi(X), \Sigma_\phi(X))$ parametered by $\phi$, which takes values from $X$ and outputs a multivariable Gaussian distribution over $Z$ to approximate the posterior distribution $p_\theta(z|x)$ [7, 11], by minimizing the Kullback-Leibler divergence $D_{KL}$ between $q_\phi(z|x)$ and the multivariable normal prior distribution $p(z) \sim \mathcal{N}(z|0, 1)$. The logarithm of $p_\theta(x|z)$ term is approximated by the expectation $\mathbb{E}_{q_\phi(z|x)}[\log p_\theta(x|z)]$. Hereby, the VAE uses the loss function called Evidence Lower BOund ($\mathcal{L}_{ELBO}$) to maximize $\log p_\theta(x)$ [7]:

$$\text{log}p_\theta(x) \geq \mathcal{L}_{ELBO} = \mathbb{E}_{q_\phi(z|x)}[\log p_\theta(x|z)] - D_{KL}(q_\phi(z|x)||p(z)) \tag{S2}$$

where the encoder $q_\phi(z|x)$ is learned by taking $X$ and optimizing $\phi$, and the decoder $p_\theta(\hat{x}|z)$ is learned by taking $Z$ and optimizing $\theta$.

Both the encoder and decoder are implemented as fully connected feedforward artificial neural networks with three hidden layers (En1, En2, En3 for encoder and De1, De2, De3 for decoder). Two Dropout layers ($p = 0.7$) are between (En1, En2) and (De2, De3). Three Batchnorm layers are between (En2, En3); (De1, De2), and De3 and the output layer. The number of units in each hidden layer is 1.5 times length of the one-hot sequence. The activation functions between linear layers is tanh, while final decoder layer uses softmax neurons. We used PyTorch [15] to implement our VAE model and trained our model using ADAM optimizer [16] with a learning rate of 0.001. We used a 3D latent space based on results of five-fold cross validation [17] by taking into consideration both validation error and gap between training and validation error. Training was conducted for 55 epochs where validation loss stopped decreasing. Codes for the vanilla VAE model are available at https://github.com/ranganathanlab/VAEforDesign.

Global sampling from the trained vanilla VAE model was conducted by randomly sampling 400 latent vectors from the Gaussian prior $p(z) \sim \mathcal{N}(0, 1)$. We passed each latent vector $z$ through the trained neural network decoder $p_\theta(x|z)$ to convert these into complete protein sequence with amino acid labels. Decoding requires multinomial sampling over the decoded probability distributions over the amino acids at each position in order to collapse the probability distribution into an unambiguous amino acid label. As such, we perform the decoding operation 10 times for each latent vector $z$ to generate a total of 4000 globally-designed sequences. Local sampling was conducted by randomly sampling 150 latent vectors from the Gaussian prior $p(z) \sim \mathcal{N}(\mu_{top}, \Sigma_{top})$, where $\mu_{top}$ and $\Sigma_{top}$ are mean and variance respectively of latent vectors of the high-r.e. natural homologs that rescue osmosensing function. For each vector, 10 sequences were generated by multinomial sampling from the decoded vector for a total of 1500 locally-designed sequences. The sequences were then filtered to eliminate highly similar sequence resulting in the production of 3984 globally-sampled sequences and 896 locally-sampled sequences (see "Gene Construction" section below).

InfoVAE inference and sequence generation

Even though VAEs have shown success across multiple domains and fields, one limitation of this modeling approach is optimizing the evidence lower bound objective (ELBO), which is prone to learning a poor amortized inference distribution $q_\phi(\mathbf{z}|\mathbf{x})$ that may not closely approximate the true and expected posterior distribution $p_\theta(\mathbf{z}|\mathbf{x})$ [18]. There are two main reasons why these issues arise: (1) inherent properties of the ELBO objective and (2) implicit modeling bias. One way to overcome these issues is to define a new training objective which learns a model to correctly reconstruct sequence and amortized inference distributions [18]. First, we used an equivalent formation of the vanilla VAE ELBO objective:

$$\mathcal{L}_{ELBO} = -\mathcal{D}_{KL}\Big(q_\theta(z)\Big|\Big|p(z)\Big) - E_{q_\phi(z)}\Big[\mathcal{D}_{KL}\Big(q_\phi(x|z)\Big|\Big|p_\theta(x|z)\Big)\Big]$$

where $D_{KL}$ is the Kullback-Leibler divergence. We include a $\lambda$ prefactor which counteracts the imbalance in terms of dimensionality of the sequence space $\mathcal{X}$ and latent space $\mathcal{Z}$. For example, in our implementation, we have $\mathbf{x} \in \mathcal{R}^{59\times21}$ and $\mathbf{z} \in \mathcal{R}^3$. To achieve an Information Maximizing VAE (InfoVAE), we will add a mutual information term $\mathcal{I}_q(x; z)$ so that the above equation becomes:

$$\mathcal{L}_{InfoVAE} = -\lambda\mathcal{D}_{KL}\Big(q_\phi(z)\Big|\Big|p(z)\Big) - E_{q_\phi(z)}\Big[\mathcal{D}_{KL}\big(q_\phi(x|z)\Big|\Big|p_\theta(x|z)\big)\Big] + \alpha\mathcal{I}_q(x; z)$$

where $\mathcal{I}_q(x; z)$ and $\alpha$ encourages the model to use the latent codes, potentially avoiding posterior collapse, and weighing the influence of this mutual information term accordingly. Since the above $\mathcal{L}_{InfoVAE}$ expression cannot be directly optimized, we can rewrite it into an equivalent form which can be optimized. By using the following definitions $\mathcal{I}_q(x;z) = E_{q_\phi(x,z)}\Big[log\frac{q_\phi(x,z)}{q_\phi(x)q_\phi(z)}\Big] = -E_{q_\phi(x,z)}\Big[log\frac{q_\phi(z)}{q_\phi(z|x)}\Big]$ and the fact that $q_\phi(x|z) = p_\mathcal{D}(x)q_\phi(z|x)/q_\phi(z)$, we can rewrite the objective as follows:

$$\mathcal{L}_{InfoVAE} = E_{q_\phi(x,z)}\Big[-\lambda log\frac{q_\phi(z)}{p_\theta(z)} - log\frac{q_\phi(x|z)}{p_\theta(x|z)} - \alpha log\frac{q_\phi(z)}{q_\phi(z|x)}\Big]$$

$$= E_{q_\phi(x,z)}\Big[logp_\theta(x|z) - log\frac{q_\phi(z)^{\lambda+\alpha-1}p_{\mathcal{D}(x)}}{p_\theta(z)^\lambda q_\phi(z|x)^{\alpha-1}}\Big]$$

$$\begin{aligned}\mathcal{L}_{InfoVAE} = E_{P_D(x)}E_{q_\phi(z|x)}\Big[log\big(p_\theta(x|z)\big)\Big] - \big(1-\alpha\big)E_{P_D(x)}\Big[\mathcal{D}_{KL}\Big(q_\phi(z|x)\Big|\Big|p_\theta(z)\Big)\Big] - \\ (\alpha+\lambda-1)\mathcal{D}_{KL}\Big(q_\phi(z)\Big|\Big|p_\theta(z)\Big) - E_{P_D(x)}\Big[log\Big(p_D(x)\Big)\Big]\end{aligned} \tag{S3}$$

where $E_{P_\mathcal{D}(x)}\Big[log\Big(p_D(x)\Big)\Big]$ is a constant with no trainable parameters that can be omitted since it does not play a role in terms of the loss gradient $\nabla\mathcal{L}_{InfoVAE}$. For our implementation, we find setting the hyperparameters $\alpha = 1$ and $\lambda = 2$ perform quite well in terms of sequence reconstruction and novel design generation. Thus, the overall expression becomes:

$$\mathcal{L} = E_{P_D(x)}E_{q_\phi(z|x)}\Big[log\Big(p_\theta(x|z)\Big)\Big] + 2\mathcal{D}_{KL}\Big(q_\phi(z)||p(z)\Big) = \mathcal{L}_{Recon} + 2\mathcal{L}_{KL} \tag{S4}$$

Furthermore, we can swap out the KL-divergence loss with a strict divergence loss, in particular the max-mean discrepancy which quantifies the distance between two distributions by comparing all of their moments when implementing the kernel embedding trick with a characteristic kernel [19–21]. Thus, the regularized term $\mathcal{L}_{KL}$ is replaced with the following expression:

$$\mathcal{L}_{MMD} = \mathcal{D}_{MMD}\Big(q_\phi(z)|p(z)\Big) = E_{p(z),p(z')}\big[k(z,z')\big] - 2E_{q(z),p(z')}\big[k(z,z')\big] + E_{q(z),q(z')}\big[k(z,z')\big]$$

where $k(\cdot,\cdot)$ is a positive definite kernel and $D_{MMD} = 0$ if and only if $p(z) = q(z)$. We choose the radial basis function (i.e., Gaussian) kernel $k(z,z') = e^{(z-z')^2/\sigma^2}$ as our characteristic kernel $k(\cdot,\cdot)$. We found that setting $\sigma$ equal to the size of the latent space led to adequate performance in learning a continuous latent space, leading to excellent generative performance via sampling $\mathbf{z}$ vectors and decoding protein sequences $\mathbf{x}$.

Both the encoder and decoder are implemented as fully connected feedforward artificial neural networks with three hidden layers (En1, En2, En3 for encoder and De1, De2, De3 for decoder). Two Dropout layers are employed between (En1, En2) and (De2, De3) with dropout hyperparameters of $p = 0.3$ and $p = 0.7$. The number of units in each hidden layer is 1.5 times length of the one-hot encoded sequence ($59 \times 21 = 1239$). The activation function between linear layers along the encoder is leaky ReLU with 0.1 negative slope hyperparameter, while the activation function is simple ReLU functions between linear layers along the decoder. The final activation function for the decoder is a softmax function, which maps the logits to categorical probability distributions for each amino acid position along the whole sequence. We used Tensorflow [22] and Keras [23] to implement our MMD-InfoVAE model and trained our model using ADAM optimizer [16] with a learning rate of 0.0001. We used a 3D latent space based on results of five-fold cross validation [17] by taking into consideration both validation error and gap between training and validation error. Training was conducted for 1000 epochs with batch size equal to 128 where validation loss stopped decreasing. Codes for the MMD-InfoVAE model are available at https://github.com/PraljakReps/InfoVAE-SH3_orthology.

Global sampling from the trained MMD-InfoVAE model was conducted by randomly sampling 2000 latent vectors from the Gaussian prior $p(z)$, $z \sim \mathcal{N}(0, I)$. We passed each latent vector z through the trained neural network decoder $p_\theta(x|z)$ to convert these into complete protein sequence with amino acid labels. We converted and decoded probabilities along each amino acid position to amino acid labels by using the argmax function, which assigns the amino acid based on the highest probability label. Local sampling was conducted by randomly sampling 1000 latent embeddings from an anisotropic Gaussian distribution estimated by the functional Sho1 embedded orthologs in the 3D latent space. The sequences were then filtered to eliminate highly similar sequence resulting in the production of 2000 globally-sampled sequences and 987 locally-sampled sequences (see "Gene Construction" section below).

Convex hull analysis

To inspect relationship between location in the latent space and functionality, convex hull analysis was performed through the scipy.spatial.ConvexHull method [24] with a tolerance of $10^{-12}$. The hull in the latent space was defined by all functional sequence (r.e. $> 0.5$) in the whole dataset of 5299 sequences. For outlier removal, since these functional sequences do not form any well-defined distribution, we excluded sample points with latent coordinate $z_0 < (-0.4)$ or $z_1 > 0.0$.

Calculation of Kullback-Leibler relative entropy $D$

Computation of $D$ is based on our previous work of Statistical Coupling Analysis (SCA) [25]. For position $i$ in our MSA, we have

$$D_i = \sum_{a=0}^{20} f_i^a \ln \frac{f_i^a}{\bar{q}^a} \tag{S5}$$

where $\bar{q}^a = (1 - \bar{q}^0) q^a$, $\bar{q}^0$ represents the fraction of gaps in the alignment, and $q^a$ is the background distribution of amino acid $a$ computed over the non-redundant database of protein sequences. $f_i^a$ is the observed frequency of amino acid $a$ at position $i$ in the MSA where length of each sequence is 59.

Gene construction

Before gene construction, 1-3 positions of a small number of designed sequences were hand-adjusted to correct effect of misalignment in the training data. Residues 16D, 17D and 46A of Sho1 (PDB 2VKN) were inserted into each designed sequence to make a final length of 62. To avoid over-similarity, sequence samples were successively picked and filtered to maintain at least 3 amino acids distance away from any other candidates in each sample set.

*S. cerevisiae* codon-optimized genes coding for all synthetic SH3 proteins were amplified from a mixed pool of oligonucleotide fragments synthesized on microarray chips (Twist). The oligonucleotides corresponding to each gene were designed with primer annealing sites and a padding sequence to make them uniform 300-mer. PCR was performed using KAPA-Hifi polymerase with 1X KAPA HiFi Buffer (Roche), 0.2 mM dNTPs and 1.0 $\mu$m of each forward (5'-CCGGTTGTACCTATCGAGTG-3') and reverse primer (5'-GACCATGCAAGGAGAGGTAC-3') in 25 $\mu$l total volume, with an initial activation (95°C, 2 min), followed by 14 cycles of denaturation (95°C, 20 s), annealing (65°C, 10 s) and primer extension (70°C, 10 s). A final extension step (70°C, 2 min) was performed subsequently. Amplified products were column purified (Zymo Research), digested with EcoR1 and BamH1, ligated into the digested PRS136 plasmid with N-terminal membrane domain of Sho1 [26], and transformed into Agilent Electrocompetent XL1-Blues to yield >250× transformants per gene. The entire transformation was cultured in 50 ml LB media containing 100 $\mu$g/ml sodium ampicillin (Amp) at 37°C overnight after which plasmids were purified and pooled.

Yeast transformation

The haploid *S. cerevisiae* strain SS101 was constructed on the W303 background gifted by Wendell Lim (UCSF) [26]. Genetic knockouts of *Ssk2* and *Ssk22* were created to remove the Sho1-independent branch of the osmoresponse pathway [27]. The pooled pRS316 plasmids with the SH3 gene library were transformed into SS101 cells using the LiAc-PEG high efficiency transformation protocol [28]. Plate check was performed to confirm at least 50 copies of each gene were successfully transformed. Transformed SS101 cells were grown in liquid Sc-Ura media for 24 h (add 20 mL Sc-Ura media for each $10^8$ total transformed cells) at 30°C, and then passaged to 250 mL fresh liquid Sc-Ura media to make OD = 0.05. After another 24 h of growth at 30°C, the Sc-Ura culture can be kept at 4°C for up to two weeks.

SH3 domain selection assay

All growth was at 30°C on shaker. The stock Sc-Ura culture was transferred to YPD media for a 24 h growth to get the $t_0$ sample. The culture was diluted every 8 h to keep the cell density below 0.2 $OD_{600}$. A small volume of the $t_0$ sample was transferred to YPD media supplemented with either (1) no KCl (non-selective) or (2) 1M KCl (selective), and the rest was span down and minipreped to extract plasmids from yeast. Both non-selective and selective cultures were grew for 24 h with $OD_{600}$ maintained under 0.2 to obtain the $t_{24}$ samples. The two $t_{24}$ samples were span down and minipreped using the same protocol as the $t_0$ sample.

Plasmids purified from both $t_0$ and $t_{24}$ samples were amplified using two rounds of PCR with Q5 polymerase (New England Biolabs) to add adapters and indices for Illumina sequencing. In the first round the DNA was amplified using primers that add from 6 to 9 random bases (Ns) for initial focusing, as well as part of the i5 or i7 Illumina adapters. Six cycles were used to minimize amplification-induced bias, followed by ampure purification before the second round PCR. In the second round of PCR, the remaining adapter sequence and TruSeq indices were added, where 20 cycles were used. The final products were gel purified (Zymo Research) , quantified using Qubit (ThermoFisher) and sequenced in an Illumina MiSeq system with a paired-end 300 cycle kit. Allele counts were obtained using standard procedures. Paired-end reads were joined using FLASH, trimmed to the EcoR1 and BamH1 cloning sites and translated. Only exact matches to the designed genes were counted. Enrichment (*en*) and relative enrichment (*r.e.*) values for each gene $x$ of the three growth conditions were calculated according to equation:

$$en(x) = \log_{10}\left(\frac{f^x_{t24}}{f^x_{t0}}\right) \tag{S6}$$

$$r.e.(x) = \frac{en(x) - en(null)}{en(wt) - en(null)} \tag{S7}$$

where $f^x_{t24}$ and $f^x_{t0}$ represents the frequency of observing gene $x$ in the $t_{24}$ and $t_0$ sample, respectively. The wild-type sequence (*wt*) is the Sho1 gene of *S. cerevisiae* and the *null* genes are TAGNTAATTTCGGCGTGGGTATGGTGGCAGGCCCCGTGGCCGGGG GACTGTTGGGCGCCATCTCCTTGCATGCACCATTCCTTGCGGCGGCGGTGCT CAACGGCCTCAACCTACTACTGGGCTGCTTCCTAATGCAGGAGTCGCATAAG GGAGAGCGTCGAGAT, where the stop codon TAG produces a Sho1 without the C-terminal SH3 domain. A second independent selection assays was performed to ensure reproducibility (Fig. S4A). The average *en* of the two trials was used to calculate *r.e.*. r.e. values of SH3 variants with at least five counts in the input population in both trials were used for analysis. The r.e. scores between the natural and synthetic libraries were globally normalized to a single null allele by linear regression fitting over the 393 sequences shared between the two pools ($\rho_{\text{Pearson}} = 0.94$, $n = 393$). The standard curve relating r.e. to binding affinity was made from a set of 11 variants of *S. cerevisiae* $\text{Sho1}^{SH3}$, comprising D18I, R38L, Y56I, Y56F, Y56M, Y56A (from published data [29]), A8V, Y10M, P11D, E21G and wild-type.

Peptide synthesis

The pbs2 MAPKK peptides were synthesized with standard 9-fluorenylmethoxycarbonyl (Fmoc) chemistry in Protein Chemistry Technology Center of UT Southwestern Medical

Center. Molecular masses were verified by mass spectrometry. Concentrations were verified by quantitative amino acid analysis.

Protein expression and purification

pET-28b plasmids encoding selected C-terminally $His_6$-tagged versions of functional SH3 domains were transformed into *E. coli* strain BL-21 (DE3). 1L of TB media containing 50 $\mu$g/mL kanamycin were inoculated 1:1000 with overnight starter LB cultures, grown in 37°C and 200 rpm to an $OD_{600}$ of 0.8-1.2, induced with 200 $\mu$M IPTG and further incubated at 18°C overnight. Cells were harvested by centrifugation (2560 g, 15min) and resuspended in 500 mM NaCl, 10 mM imidazole, 25 mM Tris-HCl, pH 8.0 and 1:1000 Tween® 20 detergent, lysed by sonication on ice (3 rounds for each 100 mL cell suspension, 90% amplititude, 2s on 2s off for 1 min in total) with 1mM PMSF, 10 $\mu$g/mL leupeptin and 2 $\mu$g/mL pepstatin, and centrifuged at 48000 g for 1 h at 4°C. SH3 proteins were purified from the cleared lysate by Ni-NTA affinity chromatography (Qiagen) , dialyzed overnight in 100 mM NaCl, 50 mM Tris, pH 7.5, and run through size exclusion in fast protein liquid chromatography (AKTA Pure 25 L1). Purified SH3 protein can be flash frozen and stored at (-80)°C.

Biophysical evaluation of SH3 *in vitro* binding assay

Binding affinity to synthetic pbs2 MAPKK ligands for $Sho1^{SH3}$ domains were measured by increase in the intrinsic tryptophan fluorescence on titration of peptide ligand into a solution of $Sho1^{SH3}$ protein at a fixed concentration of 0.25 $\mu$M (less than one fourth of expected $K_d$) in HEPES buffer (20mM HEPES, 50mM NaCl, pH 7.3-7.6). Fluorescence titration was performed on Fluorolog-3 with $\lambda_{ex} = 296$ nm and $\lambda_{em} = 330$ nm. Data were fitted to the equation

$$y = F_{min} + (F_{max} - F_{min}) \left( \frac{x}{K_d + x} \right) \tag{S8}$$

with scipy.optimize.curve_fit module in Python, where $y$ is the fluorescence reading, $x$ is ligand concentration, $K_d$ is dissociation constant, $F_{min}$ and $F_{max}$ are minimum and maximum fluorescence values.

Melting temperature ($T_m$) measurements

The $T_m$ of SH3 domains was determined using a MicroCal VP-Capillary DSC (Malvern Instruments). Purified protein (60 $\mu$M-180 $\mu$M) in 100mM NaCl, 50mM TrisHCl, pH 7.5 was heated from 10°C-110°C at 60°C/hour and the resulting curve was fit using a two-state model.

[6] Sam Sinai, Eric Kelsic, George M Church, and Martin A Nowak. Variational autoencoding of protein sequences. *arXiv preprint arXiv:1712.03346*, 2017.

[7] Carl Doersch. Tutorial on variational autoencoders. *arXiv preprint arXiv:1606.05908*, 2016.

[8] Xiaojie Guo, Sivani Tadepalli, Liang Zhao, and Amarda Shehu. Generating tertiary protein structures via an interpretative variational autoencoder. *arXiv preprint arXiv:2004.07119*, 2020.

[14] Sam Sinai, Nina Jain, George M Church, and Eric D Kelsic. Generative AAV capsid diversification by latent interpolation. *bioRxiv*, page 2021.04.16.440236, 2021.

[15] Adam Paszke, Sam Gross, Soumith Chintala, Gregory Chanan, Edward Yang, Zachary DeVito, Zeming Lin, Alban Desmaison, Luca Antiga, and Adam Lerer. Automatic differentiation in pytorch. In *31st Conference on Neural Information Processing Systems (NIPS 2017)*, 2017.

[16] Diederik P Kingma and Jimmy Ba. Adam: A method for stochastic optimization. *arXiv preprint arXiv:1412.6980*, 2014.

[17] Ron Kohavi. A study of cross-validation and bootstrap for accuracy estimation and model selection. In *International Joint Conference on Artificial Intelligence (IJCAI)*, volume 14, pages 1137–1145, 1995.

[18] Shengjia Zhao, Jiaming Song, and Stefano Ermon. Infovae: Balancing learning and inference in variational autoencoders. In *Proceedings of the AAAI Conference on Artificial Intelligence*, volume 33, pages 5885–5892, 2019.

[19] Arthur Gretton, Karsten Borgwardt, Malte Rasch, Bernhard Schölkopf, and Alex Smola. A kernel method for the two-sample-problem. *Advances in Neural Information Processing Systems*, 19:513–520, 2006.

[20] Yujia Li, Kevin Swersky, and Rich Zemel. Generative moment matching networks. In Francis Bach and David Blei, editors, *Proceedings of the 32nd International Conference on Machine Learning*, volume 37 of *Proceedings of Machine Learning Research*, pages 1718–1727, Lille, France, 07–09 Jul 2015. PMLR.

[21] Gintare Karolina Dziugaite, Daniel M Roy, and Zoubin Ghahramani. Training generative neural networks via maximum mean discrepancy optimization. *arXiv preprint arXiv:1505.03906*, 2015.

[22] Martín Abadi, Ashish Agarwal, Paul Barham, Eugene Brevdo, Zhifeng Chen, Craig Citro, Gregory S. Corrado, Andy Davis, Jeffrey Dean, Matthieu Devin, Sanjay Ghemawat, Ian J. Goodfellow, Andrew Harp, Geoffrey Irving, Michael Isard, Yangqing Jia, Rafal Józefowicz, Lukasz Kaiser, Manjunath Kudlur, Josh Levenberg, Dan Mané, Rajat Monga, Sherry Moore, Derek Gordon Murray, Chris Olah, Mike Schuster, Jonathon Shlens, Benoit Steiner, Ilya Sutskever, Kunal Talwar, Paul A. Tucker, Vincent Vanhoucke, Vijay Vasudevan, Fernanda B. Viégas, Oriol Vinyals, Pete Warden, Martin Wattenberg, Martin Wicke, Yuan Yu, and Xiaoqiang Zheng. Tensorflow: Large-scale machine learning on heterogeneous distributed systems. *arXiv preprint arXiv:1603.04467*, 2016.

[23] François Chollet. Keras: The python deep learning library. Astrophysics Source Code Library (ascl-1806), 2018.

[24] Pauli Virtanen, Ralf Gommers, Travis E. Oliphant, Matt Haberland, Tyler Reddy, David Cournapeau, Evgeni Burovski, Pearu Peterson, Warren Weckesser, Jonathan Bright, Stéfan J. van der Walt, Matthew Brett, Joshua Wilson, K. Jarrod Millman, Nikolay Mayorov, Andrew R. J. Nelson, Eric Jones, Robert Kern, Eric Larson, C J Carey, İlhan Polat, Yu Feng, Eric W. Moore, Jake VanderPlas, Denis Laxalde, Josef Perktold, Robert Cimrman, Ian Henriksen, E. A. Quintero, Charles R. Harris, Anne M. Archibald, Antônio H. Ribeiro, Fabian Pedregosa, Paul van Mulbregt, and SciPy 1.0 Contributors. SciPy 1.0: Fundamental algorithms for scientific computing in python. *Nature Methods*, 17:261–272, 2020.
